# Membrane Curvature Promotes ER-PM Contact Formation via Junctophilin-EHD Interactions

**DOI:** 10.1101/2024.06.29.601287

**Authors:** Yang Yang, Luis A. Valencia, Chih-Hao Lu, Melissa L. Nakamoto, Ching-Ting Tsai, Chun Liu, Huaxiao Yang, Wei Zhang, Zeinab Jahed, Wan-Ru Lee, Francesca Santoro, Jen Liou, Joseph C. Wu, Bianxiao Cui

**Author notes:** Correspondence should be addressed to B.C.

## Abstract

Contact sites between the endoplasmic reticulum (ER) and the plasma membrane (PM) play a crucial role in governing calcium regulation and lipid homeostasis. Despite their significance, the factors regulating their spatial distribution on the PM remain elusive. Inspired by observations in cardiomyocytes, where ER-PM contact sites concentrate on tubular PM invaginations known as transverse tubules (T-tubules), we hypothesize that the PM curvature plays a role in ER-PM contact formation. Through precise control of PM invaginations, we show that PM curvatures locally induce the formation of ER-PM contacts in cardiomyocytes. Intriguingly, the junctophilin family of ER-PM tethering proteins, specifically expressed in excitable cells, is the key player in this process, while the ubiquitously expressed extended synaptotagmin 2 does not show a preference for PM curvature. At the mechanistic level, we find that the low complexity region (LCR) and the MORN motifs of junctophilins can independently bind to the PM, but both the LCR and MORN motifs are required for targeting PM curvatures. By examining the junctophilin interactome, we identify a family of curvature-sensing proteins, Eps15-homology domain containing proteins (EHDs), that interact with the MORN_LCR motifs and facilitate junctophilins’ preferential tethering to curved PM. These findings highlight the pivotal role of PM curvature in the formation of ER-PM contacts in cardiomyocytes and unveil a novel mechanism for the spatial regulation of ER-PM contacts through PM curvature modulation.

## Introduction

In eukaryotic cells, the endoplasmic reticulum (ER) plays a central role in membrane protein synthesis, lipid production, and calcium storage. At certain locations, the ER membrane and the plasma membrane (PM) are brought into close proximity by ER-PM tethering proteins, typically within a range of 10-30 nm^1,2^, to form ER-PM contacts. These contact sites play pivotal roles in lipid exchange, calcium signaling, and phospholipid signaling^3–5^. Disruptions in ER-PM contacts and mutations in ER-PM tethering proteins have been associated with a variety of diseases, including cardiovascular and neurodegenerative diseases ^6–10^.

Notably, ER-PM contact sites are not uniformly distributed on the PM. For example, in pancreatic acinar cells^11^ and hepatocytes^12^, ER-PM contacts are enriched at the basal membrane but are nearly absent from the apical region. T cells preferentially form ER-PM contacts at immunological synapses^13^, while neurons form dense ER-PM contacts in dendrites and sparse contacts in axons ^14,15^. The spatial organization of ER-PM contacts is believed to function as a mechanism regulating local calcium influx and subcellular responses. However, the precise mechanisms governing the spatial organization of ER-PM contacts on the PM remain to be fully elucidated.

An intriguing example of nonuniform ER-PM contact sites is in striated muscle cells. In these cells, ER-PM contacts preferentially form on tubular PM invaginations known as transverse-tubules (T-tubules)^16–18^, which penetrate into the cytoplasmic domain and establish extensive contacts with the ER, referred to as dyad junctions in cardiomyocytes. Early studies showed that ER-PM contacts are 5 times more likely to form on the sarcolemma, namely the PM in muscle cells, at the T-tubules compared to the sarcolemma in other areas of mammalian ventricular cardiomyocytes ^17^. Dyad junctions in cardiomyocytes are crucial for regulating the rapid influx of calcium and mediating excitation-contraction coupling. Loss of T-tubules is accompanied by disorganized dyad junctions, disrupted calcium responses, increased susceptibility to arrhythmia, and impaired contractile function of cardiomyocytes in patients with heart diseases^19^. However, the molecular mechanism underlying the enrichment of ER-PM contact at T-tubules remains largely underexplored.

The curvature of the plasma membrane is emerging as a pivotal regulator of cellular activities. Cells respond to PM curvatures through specialized proteins known as curvature-sensing proteins, which have distinct structures for sensing and influencing membrane bending^20^. Recent studies have revealed that PM curvatures actively participate in a diverse range of cellular processes, including ion channel activity^21^, membrane trafficking^22^, signal transduction^23^, and mechanotransduction^24^. In this study, we hypothesize that PM morphology can regulate the formation of ER-PM contacts through the induction of local PM curvatures and the involvement of curvature-sensing proteins. To investigate this, we employed vertical nanostructures to induce precisely controlled PM curvatures. Our study reveals that, in cardiomyocytes, PM curvature promotes the site-specific formation of ER-PM contacts, a process mediated by junctophilin 2 (JPH2), an ER-PM tethering protein. Furthermore, we find that JPH-mediated ER-PM contacts also exhibit a preference for PM curvature in non-muscle cells. On the other hand, extended synaptotagmins (E-Syts), another family of ER-PM tethering proteins, do not exhibit a preference for PM curvature. Mechanistically, our investigation identified the Eps15-homology domain containing proteins (EHDs), a crucial family of curvature-sensing proteins, that interact with JPHs and convey the preference for PM curvature.

## Results

### 1. Nanopillar-induced PM invaginations recruit dyad components in cardiomyocytes

To assess the role of PM curvature in the formation of dyad junctions, we used human induced pluripotent stem cell-derived cardiomyocytes (iPSC-CMs). These immature iPSC-CMs do not have T-tubule structures, which develop only postnatally. T-tubules are inward-extending PM invaginations with diameters ranging from 20 nm to 450 nm, pitches (lateral distance between neighboring tubules) of 1.8-2.5 μm, and depths of 1-9 μm^25^. To induce PM invaginations resembling the shape of T-tubules, we fabricated vertical quartz nanopillars measuring 200-300 nm in diameter, 2.5 μm in pitch and 1 μm in height (**Fig. 1a**). When cells are cultured on nanopillars, their PM wraps around the nanopillars and forms inward membrane tubules (**Fig. 1b**). In this work, both the endoplasmic reticulum in non-muscle cells and the sarcoplasmic reticulum in cardiomyocytes are referred to as the ER, and both the PM in non-muscle cells and the sarcolemma in cardiomyocytes are referred to as the PM.

**Fig. 1:**
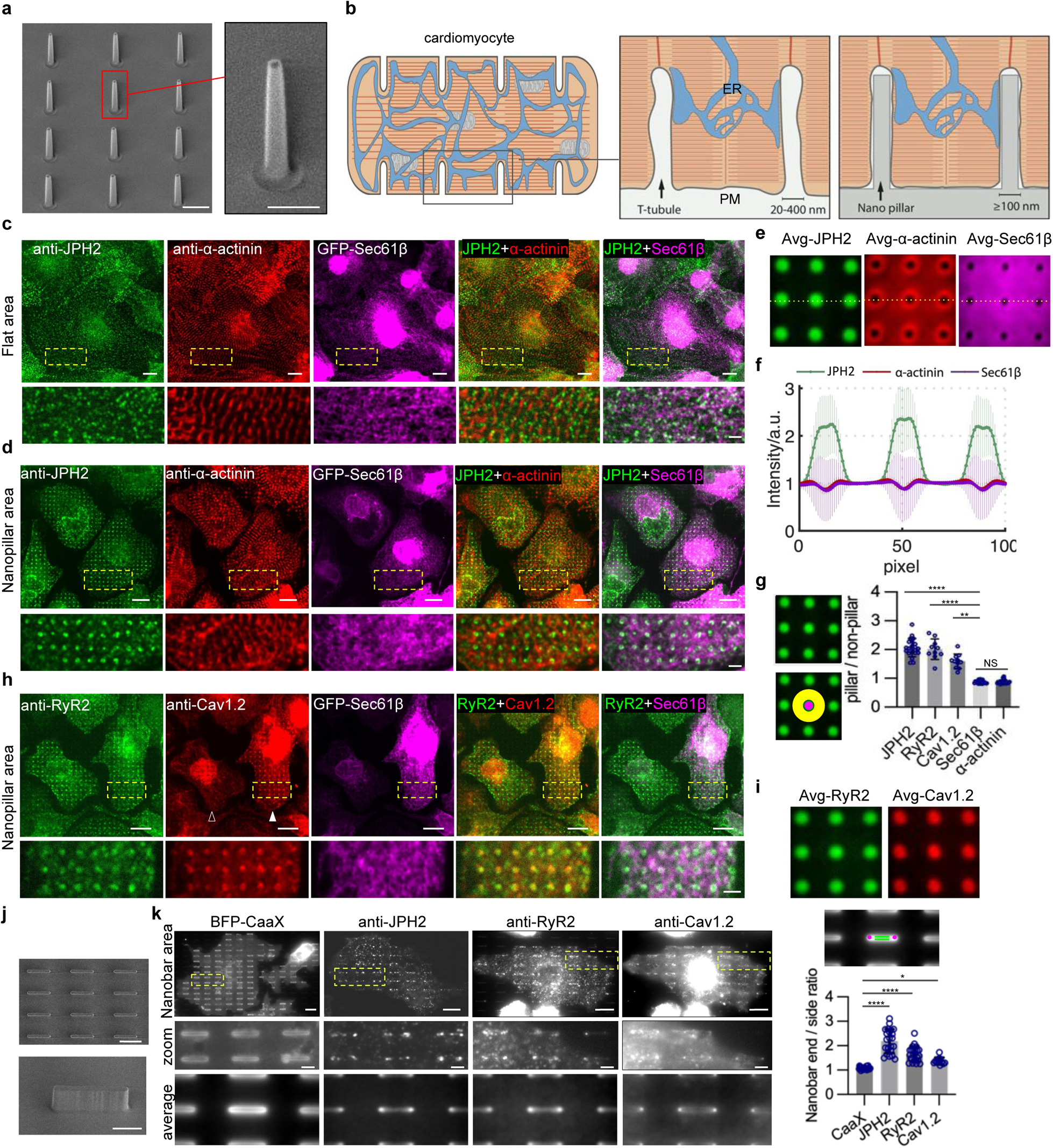
Nanopillar-induced PM invaginations recruit dyad components in cardiomyocytes. **a**, SEM images of vertical nanopillars. Scale bar 1 µm on the left, 500 nm on the right. **b**, Schematics of the T-tubule system in a cardiomyocyte (left), ER-PM contacts formed at T-tubules (middle), and nanopillar-induced membrane invaginations (right). **c-d**, Co-Immunostained JPH2 (green) and α-actinin (red) in iPSC-CMs expressing GFP-Sec61β (magenta) on flat areas (c) and on nanopillar areas (d). JPH2 (green) preferentially accumulates on nanopillar locations. Magnified images of the yellow boxes are shown at the bottom. Scale bar 10 µm on the top, 2.5 µm at the bottom. **e**, Averaged fluorescent signals for GFP-Sec61β, JPH2, and α-actinin. 101×101-pixel square regions of interest centered by each nanopillar were averaged and displayed. Over ∼3000 nanopillars were included in each condition. GFP-Sec61β n = 18 cells; JPH2 and α-actinin n = 21 cells. **f**, Intensity plots along the horizontal yellow line in e, normalized by the intensity value of the region between two nanopillars. Error bars represent standard deviation (STD) among values calculated from individual cells. **g**, The ratio of the fluorescent intensity at the nanopillars (pillar: area in the magenta mask) over the average intensity between the nanopillars (non-pillar: area inside the yellow mask). n=21, 10, 11, 18, 21 cells for JPH2, RyR2, Cav1.2, Sec61β, and α-actinin. ****P <0.0001; **P=0.0067; not-significant (NS) P>0.9999. **h**, Representative images of co-immunostained RyR2 (green) and Cav1.2 (red) in iPSC-CMs expressing GFP-Sec61β (magenta) on nanopillars. The solid arrowhead points at a cell with clear Cav1.2 signal on nanopillars and the hollow arrowhead points to a cell with minimal Cav1.2 on nanopillars. Scale bar 10 µm on the top, 2.5 µm on the bottom. **i**, Averaged fluorescent signals of ∼1200 nanopillars were included in each condition. n = 10 cells for RyR2; n = 11 cells for Cav1.2. **j**, SEM images of nanobars. Scale bar 5 µm on the top, 2 µm at the bottom. **k**, Representative images of iPSC-CMs with expressed BFP-CAAX, immunostaining of JPH2, RyR2, and Cav1.2 on nanobars. The RyR2 staining and Cav1.2 staining are the same cell. Magnified images of the yellow boxes are shown in the middle row. Averaged fluorescent signals of all nanobars from multiple cells were averaged and displayed in the bottom row. n = 23, 23, 23, 11 cells for BFP-CAAX, JPH2, RyR2, and Cav1.2. Scale bar 10 µm in the top row, 2.5 µm in the middle row. **l**, Quantification of the ratio of the fluorescent intensity at the ends of nanobars (area in the magenta mask) over the average at the sides of nanobars (area in the green mask). Cell number is the same as in (**k**). ****P <0.0001; *P=0.0277. All experiments were independently replicated at least three times. All error bars represent STD. Kruskal-Wallis test corrected with Dunn’s multiple-comparison test was used to assess significance for (**g**, **l**).

To determine whether the induction of PM invaginations by nanopillars is sufficient to induce local ER-PM contact formation in iPSC-CMs, we immunostained JPH2, a transmembrane ER protein that tethers ER membranes to the PM at contact sites in cardiomyocytes. We also co-stained α-actinin, a marker for z-lines reflecting the integrity of sarcomeres, and transfected the cells with GFP-Sec61β to visualize the general ER distribution. For cells cultured on flat areas, α-actinin staining showed a characteristic well-ordered sarcomere pattern, while JPH2 appeared as numerous small puncta widely distributed in cells (**Fig. 1c**). However, for cells cultured on nanopillar areas, JPH2 staining showed strong and preferential accumulation at regularly spaced nanopillars, in contrast to GFP-Sec61β, which did not show accumulation on nanopillars (**Fig. 1d**). Alpha-actinin staining showed that z-lines are not spatially correlated with nanopillars and largely avoided nanopillar locations. Bright-field images, PM markers, and additional representative examples of JPH2, α-actinin, and GFP-Sec61β co-imaging are included in **Extended Data Fig. 1a**.

A closer analysis of JPH2, α-actinin, and GFP-Sec61β, averaged over ∼3000 nanopillars and normalized by their intensities between nanopillars, clearly illustrated that JPH2 preferentially accumulated at nanopillar locations while α-actinin and GFP-Sec61β slightly avoided nanopillars (**Fig. 1e**). These images were generated by automatically selecting the area centered around each nanopillar and then averaging over all nanopillars in contact with cells. The intensity profiles generated along the yellow horizontal lines in Fig. 1e reveal a distinct preference of JPH2 for the nanopillars over α-actinin and GFP-Sec61β (**Fig. 1f**). We further quantified the ratio of the average fluorescence intensity at nanopillars (area within the circular magenta mask) over the intensity between nanopillars (area inside the yellow donut mask) (**Fig. 1g**). The quantification revealed a consistent ratio of ∼ 2 for JPH2 and ∼ 0.9 for both α-actinin and GFP-Sec61β (**Fig. 1g**), indicating a preferential accumulation of JPH2 at nanopillar-induced PM curvatures.

To investigate whether ER-PM contacts formed at nanopillar-induced PM invaginations incorporate crucial components of functional dyad junctions, we immunostained two key proteins, ryanodine receptor 2 (RyR2), an ER calcium release channel, and Cav1.2, a subunit of the L-type calcium channel (LTCC) present on the PM. RyR2 and LTCC are known to colocalize at dyad junctions and work in concert to facilitate voltage-induced calcium influx and subsequent store calcium release during cardiac excitation-contraction coupling^26^. We found that, similar to JPH2, RyR2 exhibited significant accumulation at nanopillar-induced PM invaginations, while GFP-Sec61β did not show notable accumulation (**Fig. 1h**). In addition, we co-stained RyR2 with Calreticulin, an ER lumen protein, as an additional general ER marker besides GFP-Sec61β, and observed a similar effect (**Extended Data Fig. 1b**). It is important to note that while all cells exhibited preferential accumulation of JPH2 and RyR2 at nanopillars, only a subset of cells exhibited such accumulation of Cav1.2 (**Fig. 1h**, filled arrowhead). In some cells, Cav1.2 appeared to be located within intracellular ER structures rather than the PM (**Fig. 1h**, empty arrowhead). This cell-to-cell variation in Cav1.2 may be attributed to the inherent immaturity and heterogeneity of iPSC-derived cardiomyocytes^27^ and the relatively late developmental expression of Cav1.2^28^. Averaged images of RyR2 and Cav1.2 (**Fig. 1i**), and quantitative analyses in cells with high Cav1.2 expression (**Fig. 1g**) demonstrated their preferential accumulation at PM invaginations induced by nanopillars, which indicates the formation of functional ER-PM contacts at PM curvatures. In addition to using iPSC-CMs generated by ourselves, we also examined iPSC-CMs from a commercial source (iCells) as well as primary rat embryonic cardiomyocytes. Immunostaining of RyR and JPH2 in these cells also showed strong accumulations at nanopillar locations (**Extended Data Fig. 1c**).

Furthermore, we engineered vertical nanobars to induce both curved and flat PMs on the same nanostructure (**Fig. 1j**). These nanobars, at 2 μm height, 200-300 nm width, 5 μm length, and 10 μm pitch, induce high PM curvatures at their vertical ends and horizontal top, and flat membranes along their sidewalls, serving as internal controls. When imaging at the mid-height of the nanobars, the PM curvature is pronounced at the nanobar ends. Staining results showed that JPH2, RyR2, and Cav1.2 all exhibit bar-end accumulations, confirming their preferential localization to membrane curvatures. In comparison, the expressed plasma membrane marker BFP-CAAX uniformly wraps around the nanobars displaying no curvature preference (**Fig. 1k, l**).

### 2. Electron microscopy and expansion microscopy reveal the promotion of ER-PM contact formation on nanopillar-induced PM invaginations

To directly visualize ER-PM contacts, we used focused ion beam (FIB) and scanning electron microscopy (SEM) to image membrane interfaces at nanopillar locations (**Fig. 2a**). We employed an ultrathin resin embedding method that enables *in situ* examination of the membrane-nanostructure interface without removing the substrate^29^. Cells cultured on nanopillar substrates were heavy metal stained to delineate membranes and then embedded in an ultrathin layer of epoxy resin to preserve cellular architectures. Cells on nanopillars were first identified under the SEM. Then we used the FIB beam to vertically mill through the sample to reveal the cell-nanopillar interface. **Fig. 2b** shows a typical SEM image of the cardiomyocyte-nanopillar interface, which includes a clear ER-PM contact formed at the left side of the nanopillar.

**Fig. 2:**
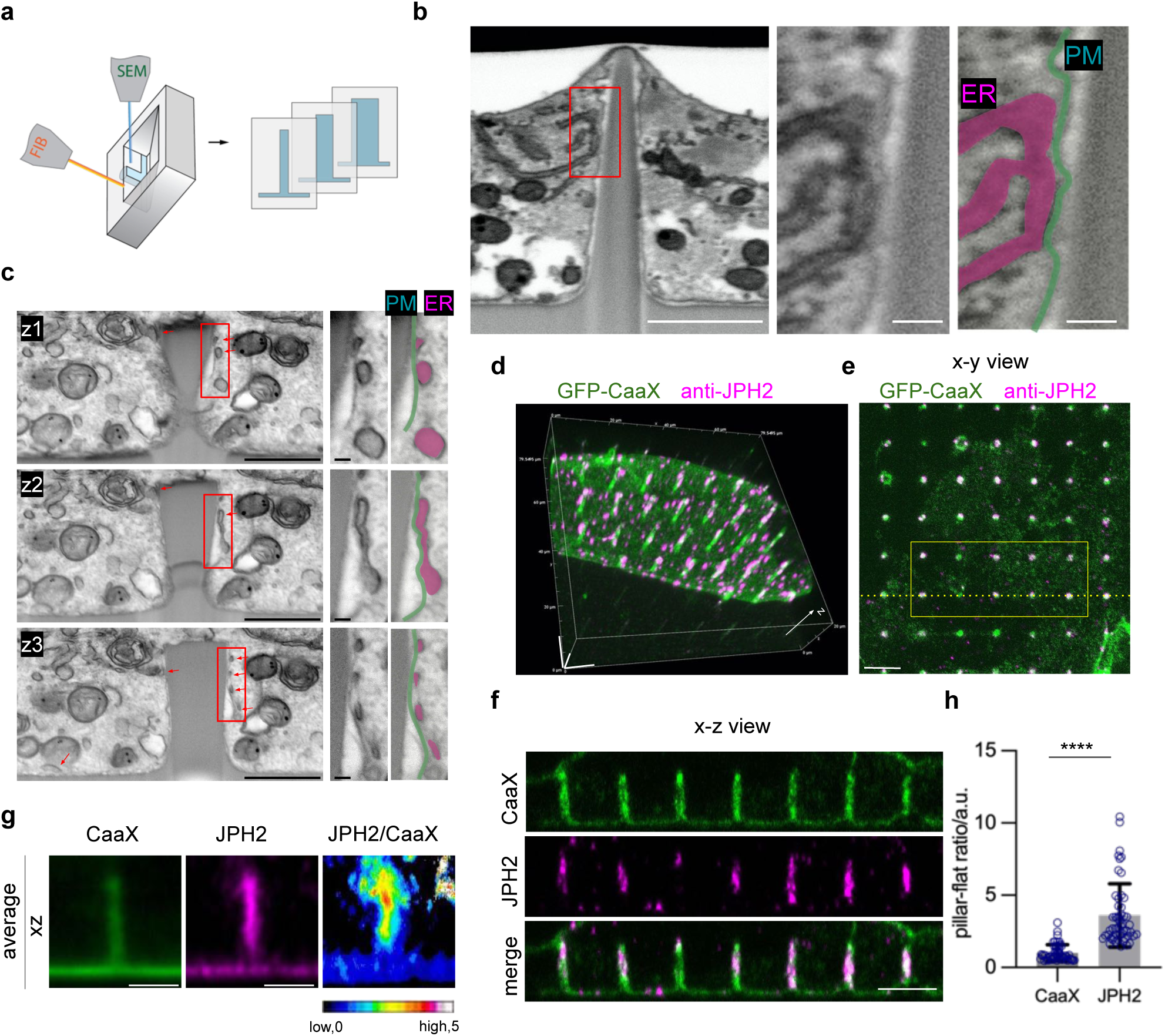
Nanopillar-induced membrane invaginations promote local formation of ER-PM contacts in cardiomyocytes. **a**, Schematic illustration of using FIB-SEM to examine the cell-nanopillar interface. **b**, A FIB-SEM image of the interface between an iPSC-CM and a nanopillar. SEM images are black-white inverted for clarity. Scale bar 1 µm. An enlarged view of the region in red box is displayed on the right side. Scale bar is 100 nm. The PM (green) and the ER (magenta) are highlighted in pseudo colors in the enlarged view. **c**, Three different FIB-SEM sections of an HL-1 cardiomyocyte on the same nanopillar. The red arrows indicate the ER-PM contact sites. Scale bars are 1 µm. Enlarged views are highlighted the same way as in (**b)**. Scale bars are 100 nm. **d**, Expansion microscopy imaging of an iPSC-CM cultured on nanopillars. GFP-CAAX in green, anti-JPH2 in magenta. Scale bars are 10 µm in each dimension. **e**, An x-y image focused on the middle height of nanopillars using expansion microscope. Scale bar 10 µm. **f,** The x-z view along the yellow dash line in (**e)**. The z-dimension was scaled by the ratio between z-step size and x-y pixel size to exhibit x and z at the same dimensional scale. Scale bar 10 µm. **g**, An averaged x-z image of 10 pillars shown in yellow box in (**e)**. The intensities were normalized by the inter-pillar intensities on flat membranes for both CAAX and JPH2 channels and then displayed at the same scale. The ratiometric image was the ratio between JPH2 and CAAX channel. Scale bar 5 µm. **h**, Quantifications of the nanopillar-to-flat intensity ratios for GFP-CAAX and JPH2, normalized by the average intensity ratio for GFP-CAAX. Each dot represents the averaged ratio for a region of nanopillars inside a cell (typically 5-6 pillars). n = 52 regions from 15 cells for each probe. ****P <0.0001. All experiments were independently replicated at least two times. All error bars represent STD. Two-tailed Mann-Whitney test was used to assess significance in (**h**).

FIB-SEM enables sequential FIB milling for volumetric SEM imaging. We obtained a series of 36 SEM images to visualize ER-PM contacts in a cardiomyocyte interfaced with a nanopillar (**Supplementary Video 1**). Multiple ER-PM contacts were observed on the PM surrounding a single nanopillar (**Fig. 2c**). The ER tubules and ER sheets are interconnected, as shown in serial images. Large ER-PM contact patches in a cross-section image can evolve into smaller, distinct patches in different cross-section images (red arrows in **Fig. 2c**). From the 36 SEM images, the length of ER-PM contacts formed on the curved PM surrounding nanopillars was ∼ 3.6 times of that formed on the flat PM of a similar length between nanopillars (**Extended Data Fig. 2**). The FIB-SEM measurements suggest that ER-PM contacts preferentially form on curved PM.

The FIB-SEM imaging method, while valuable, has limitations in terms of throughput, which restricts our ability to perform quantitative measurements. To quantitatively determine whether PM invaginations induced by nanopillars elicit preferential formation of ER-PM contacts, we used expansion microscopy (ExM)^30,31^. This technique allowed us to examine a larger number of nanopillars and cells, significantly enhancing our data robustness (**Fig. 2d, Supplementary Video 2**). By expanding our samples and thus increasing the effective spatial resolution, confocal images provided clear and distinct visualization of JPH2 puncta on the PM, both at the nanopillar sites and the flat areas. In an x-y plane image, both JPH2 and the PM marker GFP-CAAX exhibited considerably higher intensities at the nanopillar locations (**Fig. 2e**) owing to the vertical projection effect of the PM wrapping around the nanopillars. The x-z plane image shows clear preferential accumulation of JPH2 signals, compared to GFP-CAAX, on nanopillars than at flat areas between nanopillars (**Fig. 2f**).

To quantitatively assess JPH2’s preference for PM curvature, we averaged xz-plane images for a line of nanopillars and the surrounding flat area, normalizing the intensity of JPH2 against that of GFP-CAAX (**Fig. 2g**). The normalized image clearly indicated a higher density of JPH2 at curved PMs surrounding nanopillars compared to the flat area (**Fig. 2g**). Quantitative analysis of over 50 regions from 15 cells revealed that the pillar-to-flat ratio for JPH2 was 3.8 ± 1.4 times higher than the pillar-to-flat ratio of CAAX (**Fig. 2h**). Therefore, JPH2 is significantly enriched on curved PMs surrounding nanopillars compared with flat PMs of the same area, confirming that ER-PM contacts preferentially form on curved PMs in iPSC-CMs.

### 3. The JPH family of tethering proteins, but not E-Syt2, exhibits a strong preference for PM curvatures

T-tubules are usually present in specialized cells, namely, mature striated muscle cells. To examine whether JPH2-mediated ER-PM contacts also prefer PM curvatures in non-muscle cells, we transfected mCherry-JPH2 into U2OS cells, a human osteosarcoma cell line. For this study, we employed smaller nanobars, 1 μm height, 200 nm width, 2 μm length, and 5 μm pitch (**Fig. 3a**), to induce membrane curvatures in U2OS cells, as these cells are more readily deformed than CMs.

**Fig. 3:**
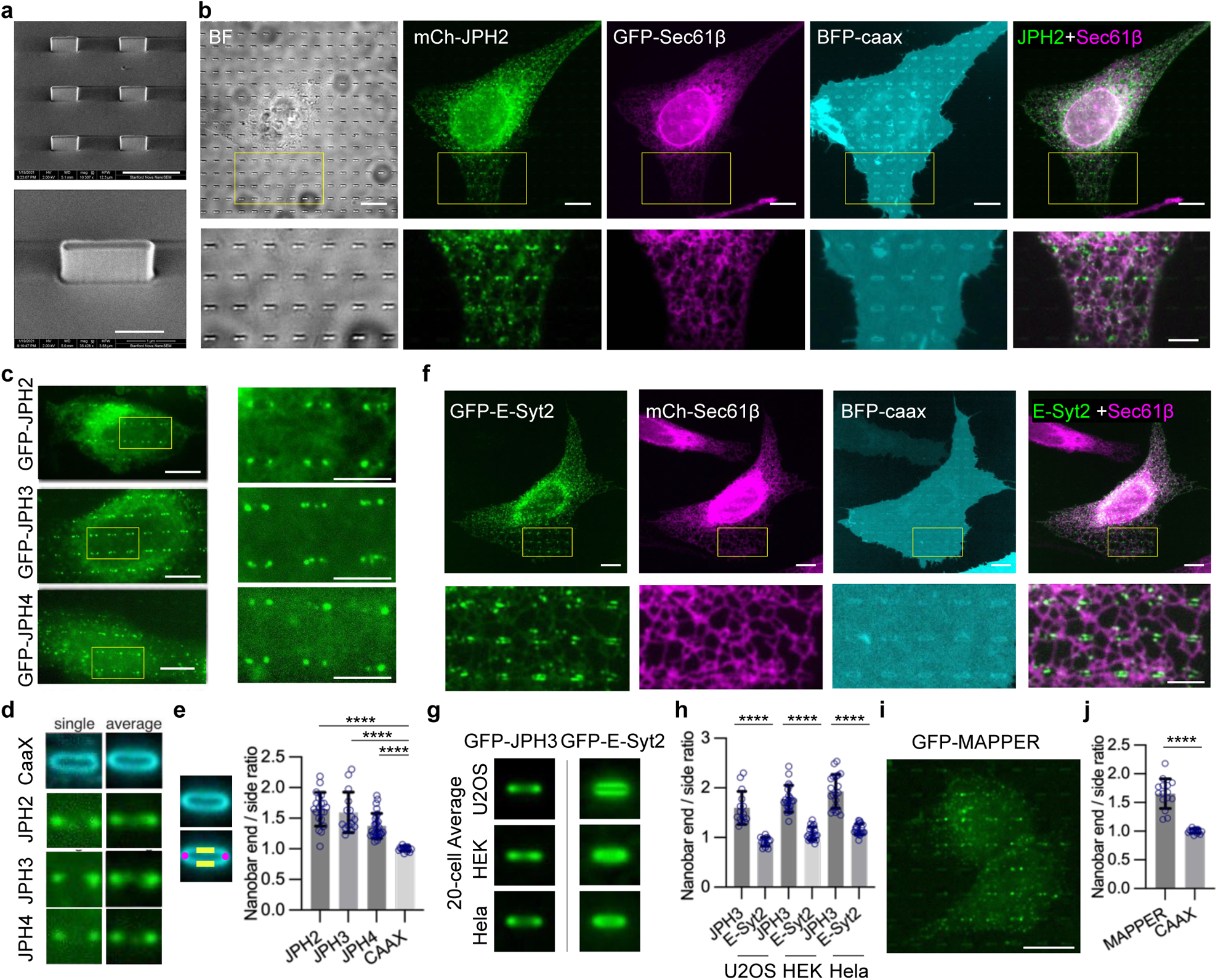
JPH family of tethering proteins, but not E-Syt2, exhibit a strong preference for PM curvatures. **a**, SEM images of nanobars. Scale bar 4 µm on the top, 1 µm at the bottom. **b**, U2OS cells co-expressing mCherry(mCh)-JPH2 (green), GFP-Sec61β (magenta), and BFP-CAAX (Cyan). mCh-JPH2 preferentially accumulates at the ends of nanobars. Bottom: Enlarged images of the regions in yellow boxes. Right: merged image of JPH2 and Sec61β. Scale bar 10 µm on the top, 5 µm at the bottom. **c**, Representative images show that GFP-JPH2, GFP-JPH3 or GFP-JPH4 preferentially accumulates at the end of nanobars. Right: Enlarged image of the yellow boxed 6 nanobars. Scale bar 10 µm on the left, 5 µm on the right. **d**, Enlarged single nanobar images and the average nanobar images of cells shown in **c**. Left: image of a single nanobar; right: the averaged image of all nanobars within a single cell. **e**, Quantification of the nanobar end-to-side ratios. The ratios are calculated using the average intensity of the bar ends (the magenta mask) divided by the average intensity of the bar sides (the yellow mask). Each dot in the quantification graph represents the average value of a single cell. n = 20, 15, 27, 15 cells for JPH2, JPH3, JPH4 and CAAX. ****P < 0.0001. **f**, U2OS cells co-expressing GFP-E-Syt2 (green), mCherry-Sec61β (magenta), and BFP-CAAX (Cyan). GFP-E-Syt2 does not preferentially accumulate at the ends of nanobars. Bottom: Enlarged images of the regions in yellow boxes. Right: merged image of GFP-E-Syt2 and mCherry-Sec61β. Scale bar 10 µm on the top, 5 µm at the bottom. **g**, Average nanobar images of GFP-JPH3 and GFP-E-Syt2 singly expressed in U2OS, HEK, and Hela cells respectively. n = 20 cells for each indicated probe in each cell line. **h**, Quantifications of the end-to-side intensity ratios for GFP-JPH3 or GFP-E-Syt2 in U2OS cell, HEK-293T cells, or Hela cells. n = 20 cells for each condition. ****P < 0.0001. **i,** U2OS cells expressing GFP-MAPPER on nanobars. Scale bar 10 µm. **j**, Quantifications of the nanobar end-to-side ratios for GFP-MAPPER and GFP-CAAX. n = 15 cells for each probe. ****P < 0.0001. All experiments were independently replicated at least three times. All error bars represent STD. Brown-Forsythe and Welch ANOVA tests was used to assess significance for (**e**,**h**). Two-tailed Welch’s t test was used to assess significance for (**j**).

In U2OS cells, mCherry-JPH2 showed selective enrichment at nanobar ends (**Fig. 3b**), indicating its preference for PM curvature. In the same cell, the ER marker GFP-Sec61β did not accumulate on the nanobars and the PM marker CAAX fused to blue fluorescence protein (BFP-CAAX) wrapped around the nanobars (**Fig. 3b**). Although JPH2 is not expressed in U2OS cells, the RNA-sequencing database shows that its homologs JPH3 and JPH4 are endogenously expressed in U2OS cells^32^. We generated GFP-JPH3 and GFP-JPH4 constructs using the U2OS cell cDNA library. When GFP-JPH2, GFP-JPH3, or GFP-JPH4 was transiently expressed in U2OS cells, all three displayed a pronounced preference for PM curvature at the nanobar ends (**Fig. 3c**). The averaged image of GFP-CAAX showed a relatively even wrapping of the PM around the nanobars (**Fig. 3d**). Meanwhile, the averaged images of GFP-JPH2, GFP-JPH3, or GFP-JPH4 revealed dumbbell distributions featuring pronounced protein accumulations at the ends of the nanobars (**Fig. 3d**). The quantifications of end-to-side intensity ratios for JPH2, JPH3, and JPH4 showed significantly higher values than that of CAAX, which has a value of ∼1 (**Fig. 3e**), indicating a strong preference of JPHs for PM curvature. We noticed that GFP-JPH2 in U2OS cells exhibited a higher ER network population compared to GFP-JPH3 and GFP-JPH4. This is likely due to U2OS cells expressing endogenous JPH3 and JPH4 but not JPH2. For subsequent investigations in U2OS cells, we selected JPH3 as the representative JPH for its lower intracellular fluorescence background.

ER-PM contacts can be mediated by several tethering protein families. To determine whether the curvature preference is unique to JPH or a shared feature of ER-PM tethers, we examined extended synaptotagmin 2 (E-Syt2), a ubiquitously expressed ER-PM tethering protein. Like JPHs, E-Syt2 is an ER-anchored protein and mediates ER-PM contact by binding to the PM through its cytosolic domain^33^. To our surprise, GFP-E-Syt2 often appeared to avoid the nanobar ends and locate along the sidewalls (**Fig. 3f**). Co-transfected mCherry-Sec61β did not accumulate on the nanobars and BFP-CAAX evenly wrapped around the nanobars (**Fig. 3f**). The averaged image of GFP-E-Syt2 illustrated protein accumulation along the two sidewalls, in sharp contrast to GFP-JPH3 accumulation at the nanobar ends (**Fig. 3g**). Furthermore, GFP-JPH3 and GFP-E-Syt2 expressed in HeLa and HEK cells showed similar behaviors (**Fig. 3g and Extended Data Fig. 3a**). Quantifications of the end-to-side ratios in the three cell lines confirm that JPH3 preferentially binds to curved PMs at the nanobar ends, whereas E-Syt2 shows no obvious preference for nanobar ends (**Fig. 3h**).

Furthermore, we investigated whether GFP-JPH3 retains its curvature preference in artificially induced ER-PM contacts using a dimerization-dependent fluorescent protein (ddFP) technology^34^. ddFP involves the reversible binding of two dark components GB and RA that form a dimeric red fluorescent protein when they are in close proximity. We constructed GB-CAAX to target the GB component to the PM and RA-Sec61β to target the RA component to the ER. At ER-PM contacts, the two components bind and become fluorescent (**Extended Data Fig. 3b**). Overexpression of GB-CAAX and RA-Sec61β induced extensive and large ER-PM contacts (**Extended Data Fig. 3c**), likely due to the relatively low dissociation rate between RA and GB^34^. Interestingly, although GFP-JPH3 entirely colocalized with ddFP in contact patches on flat areas, JPH3 showed a much more pronounced preference toward the end of the nanobars than ddFP (**Extended Data Fig. 3d**), further confirming that JPH3 exhibits a preference for curved PMs.

Finally, to directly visualize the localization of ER-PM contacts in U2OS cells without overexpressing tethering proteins, we utilized a previously developed ER-PM marker, GFP-MAPPER (membrane-attached peripheral ER), which has an ER transmembrane domain conjugated with flexible linkers and a polybasic motif targeting the PM. At low expression levels, MAPPER is incorporated into existing ER-PM contacts with minimum perturbations ^35^. We found that GFP-MAPPER exhibited an obvious curvature preference with an end-to-side ratio of 1.65 ± 0.26, suggesting that endogenous ER-PM contacts in U2OS cells preferentially form on curved PM (**Fig. 3i, j**). When co-expressed with mCherry-JPH3, MAPPER showed strong accumulation at the nanobar ends. However, when co-expressed with E-Syt2, MAPPER colocalized with E-Syt2 with no obvious curvature preference, further confirming that JPH3-mediated, but not E-Syt2-mediated, ER-PM contacts preferentially form on curved plasma membranes (**Extended Data Fig. 3e**).

### 4. STIM1 and ORAI1 are incorporated into ER-PM contacts formed on curved PM following calcium depletion

ER calcium depletion-induced calcium entry, referred to as store-operated calcium entry (SOCE), is indispensable for the homeostasis of intracellular Ca^2+^, and this process is highly dependent on two major molecular players: STIM1 and ORAI1. STIM1, an ER-resident calcium sensor, oligomerizes and translocates to ER-PM contact sites when the ER Ca^2+^ level decreases ^36,37^. At the ER-PM contact sites, STIM1 interacts with the PM-resident ORAI1 calcium channel across the junctional space to facilitate SOCE from the extracellular space^38–41^. STIM1 has been recognized to primarily localize to pre-existing ER-PM contacts upon activation^36^. In this section, we examined whether STIM1 and ORAI1 could be incorporated into ER-PM contact sites formed on curved PM upon Ca^2+^ depletion. U2OS cells were used in most of the subsequent studies because these cells are easy to transfect and exhibit clear localization of ER-PM contacts to PM curvatures.

In the resting state, GFP-JPH3 was observed to form ER-PM contacts at the nanobar ends, while mCherry-STIM1 displayed a predominantly intracellular pattern in the ER network with minimal localization to nanobars (**Fig. 4a**). Upon a 5-min treatment with 2 μM thapsigargin (Tg) to deplete the ER calcium store, mCherry-STIM1 underwent clustering at the nanobar ends, where it colocalized with GFP-JPH3 (**Fig. 4a**). ORAI1 exhibited a relatively uniform distribution around the nanobars in the resting state, typical of membrane proteins that have no curvature preference (**Fig. 4b**). Tg treatment induced dramatic redistribution of ORAI1-GFP to ER-PM contact sites located at the nanobar ends, where it colocalized with STIM1 (**Fig. 4b, Supplementary Video 3**). Within our temporal resolution of 0.1 frame/second, ORAI1 exhibited similar clustering kinetics on curved PM and flat PM (**Fig. 4c**). On curved PMs, STIM1 and ORAI1 displayed similar accumulation kinetics within our temporal resolution (**Fig. 4d**). Averaged nanobar images (**Fig. 4e**) and end-to-side ratio quantification (**Fig. 4f**) confirm that STIM1 and ORAI1 translocated to the nanobar ends upon store Ca^2+^ depletion.

**Fig. 4:**
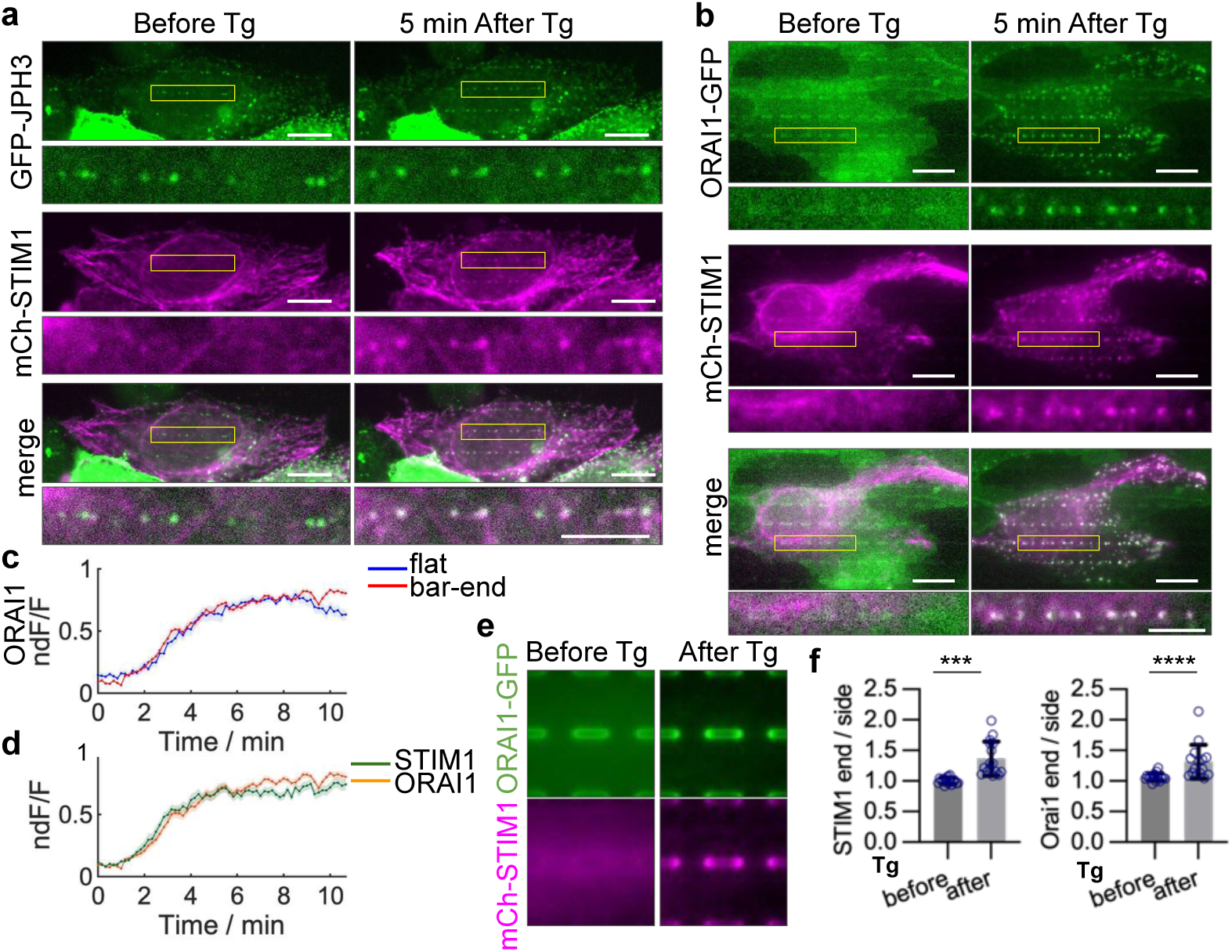
STIM1 and ORAI1 are incorporated into ER-PM contacts formed on curved PM following calcium depletion. **a**,**b**, U2OS cells co-transfected with mCherry-STIM1 and GFP-JPH3 (**a**) or ORAI-GFP (**b**) before and 5-10 min after 10 µM Tg treatment. Tg treatment induced accumulations of mCherry-STIM1 and ORAI1-GFP at nanobar ends. Enlarged views of the region in yellow boxes are displayed at the bottom of each image. Scale bar 10 µm in whole cell images, 5 µm in zoom-in images. **c**, Relative time-dependent increase (dF/F) of ORAI1 cluster intensities at the bar-end and on the flat area in cells co-transfected with mCherry-STIM1 and ORAI1-GFP upon Tg treatment. Intensity of regions of 0.85×0.85 µm^2^ at ORAI1 clusters formed at nanobar ends or on flat areas were calculated. Representative averaged dF/F traces are shown. n = 21 regions from 7 cells. Shaded error bars represent SEM. **d**, Time-dependent increases of STIM1 and ORAI1 cluster intensities at bar ends upon Tg treatment. Average intensity of STIM1 and ORAI1 were calculated from the same cells and the same regions as in (**c**). dF/F traces were normalized to their plateau value before average (ndF/F) to compare between different probes. Shaded error bars represent SEM. **e**, Averaged nanobar images for mCh-STIM1 and ORAI1-GFP, before and after the Tg treatment. N = 16 cells for each condition. **f**, Quantifications of the end/side ratios for mCh-STIM1 and ORAI1-GFP, before and after the Tg treatment. n = 16 cells for each condition. ***P = 0.0001, ****P <0.0001. All experiments were independently replicated at least three times. All error bars represent STD unless otherwise stated. Two-tailed Welch’s t test was used to assess significance for STIM1 in (**f**), and Two-tailed Mann-Whitney test was used for ORAI1 in (**f**).

### 5. A conserved polybasic sequence in the low complexity (LCR) region and the MORN motifs synergistically mediate the PM binding and the curvature sensing of JPH3

JPHs are composed of eight conserved MORN (membrane occupation and recognition nexus) motifs with a long LCR joining segment positioned between MORN6 and MORN7, followed by an α-Helix domain (αHelix or αHlx) and an ER transmembrane (TM) domain (**Fig. 5a**). Some studies have suggested that MORN repeats tether the PM by binding to phospholipids ^42,43^, while other studies have indicated weak or no phospholipid interactions for MORN motifs ^44,45^. The LCR joining region is structurally flexible without a known function. It has been suggested to act as a steric hindrance that reduces the binding affinity between JPH and LTCC^45^. Here, we engineered a series of truncations and mutations in JPH3 to identify the motif(s) responsible for JPH3’s PM binding and curvature sensing (**Fig. 5a**).

**Fig. 5:**
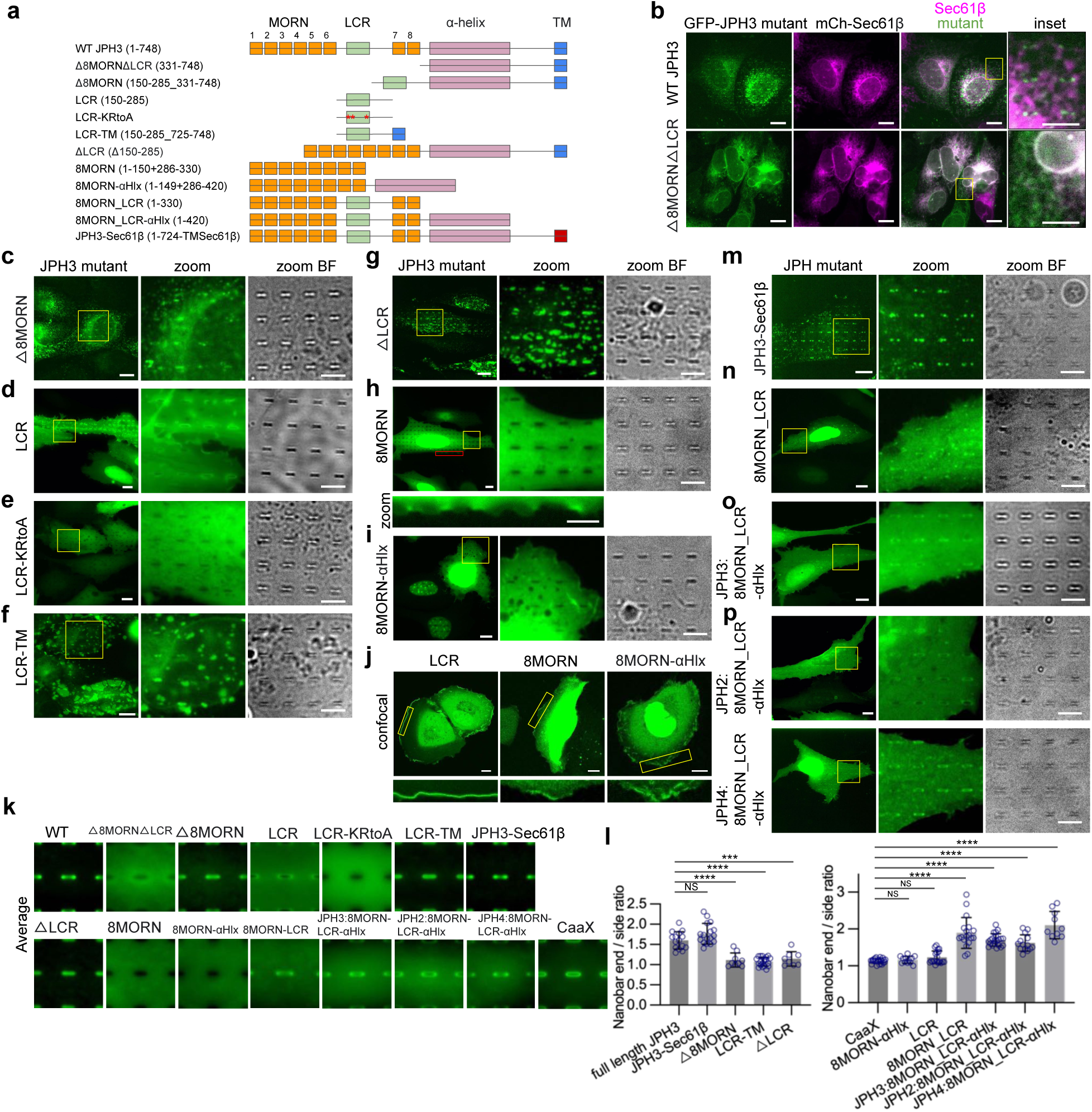
A conserved polybasic sequence LCR region and the MORN motifs synergistically mediate the PM binding and the curvature sensing of JPH3. **a**, The domain structure of JPH3 and eight engineered mutants of JPH3. The numbers in brackets refer to the amino acid positions of the residues. **b**, U2OS cells co-expressing mCherry-Sec61β with GFP-JPH3 or GFP-△8MORN△LCR. While GFP-JPH3 preferentially accumulates at nanobar ends, GFP-△8MORN△LCR is diffusive on the ER network. Inset: enlarged view of the yellow boxes. **c-f**, Representative images of U2OS cells expressing GFP-tagged JPH3 constructs: △8MORN (**c**), LCR (**d**), LCR-KRtoA (**e**), LCR-TM (**f**). Left: whole cell images; Middle: enlarged views of the regions in yellow boxes; Right: Bright field images of the same enlarged areas. **g-i**, Representative images of U2OS cells expressing GFP-tagged JPH3 constructs: △LCR (**g**), 8MORN (**h**), 8MORN-αHelix (**i**) on nanobars. Left: whole cell images; Middle: enlarged view of the region in yellow boxes; Right: the corresponding bright field images of the enlarged view. Bottom in (**h**): enlarged view of the region in the red box. **j**, Representative confocal images of U2OS cells expressing GFP-LCR, mCh-8MORN, or GFP-8MORN-αHelix. Top: whole cell images; Bottom: enlarged view of the regions in yellow boxes. **k**, Averaged nanobar images in cells transfected with the indicated probes, displayed at the same contrast scale. The image area is 10 µm x 10 µm. Cell number: n = 15 (full-length JPH3), 20 (△8MORN△LCR), 9 (△8MORN), 20 (LCR), 16 (LCR-KRtoA), 20 (LCR-TM), 17 (JPH3-Sec61β), 8 (△LCR), 20 (8MORN), 20 (8MORN-αHelix), 20 (8MORN_LCR), 21 (JPH3:8MORN_LCR-αHelix), 13 (JPH2:8MORN_LCR-αHelix), 10 (JPH4:8MORN_LCR-αHelix), and 20 (CAAX). **l**, Left: Quantifications of the nanobar end-to-side ratios for constructs with the TM domain including full-length JPH3, JPH3-Sec61β, △8MORN, LCR-TM, and △LCR; Right: Quantifications of the nanobar end-to-side ratios for constructs without TM domain including: CAAX, 8MORN-αHelix, LCR, 8MORN_LCR, 8MORN_LCR-αHelix of JPH3, of JPH2, and of JPH4. Cell number is the same as in (**k**). ****P < 0.0001, ***P=0.0002, NS (JPH3-Sec61β) P = 0.2232, NS (8MORN-αHelix) P = 0.8345, NS (LCR) P = 0.1468. **m-p**, Representative images of U2OS cells expressing GFP-tagged JPH3 constructs: JPH3-Sec61β (**m**), 8MORN_LCR (**n**), 8MORN_LCR-αHelix of JPH3 (**o**), and 8MORN_LCR-αHelix of JPH2 and JPH4 (**p**) on nanobars. Left: whole cell images; Middle: enlarged views of the region in yellow boxes; Right: the corresponding bright field images of the enlarged views. Scale bars 10 µm in all whole cell images, and 5 µm in all zoom-in images. All experiments were independently replicated at least three times. All error bars represent STD. Brown-Forsythe and Welch ANOVA test was used to assess significance for (**l**, both left and right).

We found that the Δ8MORNΔLCR construct, which is the truncation of JPH3’s N-terminal 8 MORN domains along with the LCR, entirely lost both the PM targeting and the curvature sensing capability (**Fig. 5b**). The Δ8MORNΔLCR construct colocalizes with Sec61β at the intracellular ER network. This result is consistent with previous studies showing that the N-terminal fragment is crucial for JPH3’s ER-PM tethering function ^42^.

To determine whether MORN motifs are crucial for the PM tethering, we constructed a Δ8MORN mutant by directly linking LCR to the Δ8MORNΔLCR construct. Surprisingly, unlike Δ8MORNΔLCR, Δ8MORN formed discrete ER-PM contacts (**Fig. 5c**), suggesting that the LCR segment binds directly to the PM or indirectly to components in ER-PM contacts. To test whether LCR directly binds to the PM, we fused the LCR segment to a GFP tag (GFP-LCR) without any transmembrane domain. Interestingly, GFP-LCR showed strong PM localization as well as nuclear accumulation (**Fig. 5d**), demonstrating that the LCR segment has a high affinity for the PM. After examining the LCR sequence in the 4 human JPH genes, we identified a conserved polybasic sequence in the LCR segment (**Extended Data Fig. 4a**) and hypothesized that the cationic charges are responsible for GFP-LCR localization to the PM and the nucleus. This hypothesis was confirmed by the observation that mutations of the 6 cationic amino acids (LCR-KRtoA: K210A, K211A, K212A, K224A, R226A, K227A) completely abolished both the PM targeting and the nuclear localization of GFP-LCR (**Fig. 5e**). To further examine whether the high affinity of LCR for the PM is sufficient to induce ER-PM contact formation, we directly linked LCR to the ER transmembrane domain (LCR-TM). Unlike the homogenous distribution of LCR on the PM, LCR-TM is spatially constrained by the ER network and displays large and discrete patches at ER-PM contact sites (**Fig. 5f**), confirming the capability of LCR-TM to function as an ER-PM tethering protein. These results demonstrate that the LCR segment in JPH3 tethers the PM, likely through electrostatic interactions between its polybasic sequence and negatively charged phospholipids.

Next, to investigate the role of MORN motifs in PM targeting, we generated a ΔLCR mutant (**Fig. 5a**). Surprisingly, ΔLCR is efficiently incorporated into ER-PM contacts (**Fig. 5g**), suggesting that the MORN domains either directly or indirectly bind to the PM. To confirm this, we constructed GFP-8MORN, lacking the LCR region and the ER transmembrane domain. When expressed in U2OS cells, GFP-8MORN appeared diffusive and cytosolic without obvious nanobar wrapping (**Fig. 5h**). However, membrane ruffle-like features were observed at cell edges, indicating weak PM binding (**Fig. 5h**). To investigate the potential enhancement of structural stability of MORN by the α-helical domain^45^, we generated GFP-8MORN-αHelix. We found that GFP-8MORN-αHelix is mostly nuclear localized, but it also displays clear membrane ruffles at the cell periphery, indicating PM affinity (**Fig. 5i**). To better visualize PM localization, we used confocal microscopy to image LCR, 8MORN, and 8MORN-αHelix. As shown in **Fig. 5j**, the PM localization of 8MORN-αHelix is stronger than that of 8MORN, likely due to the structural stability provided by the αHelix, but its PM affinity is much less than that of LCR.

Although Δ8MORN, ΔLCR, and LCR were all able to bind the PM, they did not exhibit obvious curvature preference, as shown in **Fig. 5c, d, g** and by the averaged nanobar images in **Fig. 5k**. It is worth noting that the behavior of each ER-PM tethering construct is greatly affected by its expression level. Specifically, when the expression level is very low, the constructs are sorted into existing contacts. Whereas when the expression level is too high, many tethering proteins induce extensive growth of ER-PM contact patches that are larger than the normal range. Therefore, we chose intermediate expressions for quantitative measurements to assess the curvature preference of specific constructs in ER-PM contacts (see Methods for details). Additionally, constructs without the TM domain typically exhibit a higher cytosolic background, elevating the baseline for the quantification and resulting in relatively lower ratios compared to those with the TM domain. Therefore, we compared the constructs with and without the TM domain separately (**Fig. 5l**). Quantitative analysis of the nanobar end-to-side ratios showed that the curvature preferences of GFP-Δ8MORN, GFP-ΔLCR, and GFP-LCR-TM were severely compromised compared to full-length JPH3 (**Fig. 5l**). Furthermore, GFP-LCR, GFP-8MORN, and GFP-8MORN-αHelix showed no obvious curvature preference compared to the membrane marker CAAX (**Fig. 5l**). These results indicate that either the LCR region or the MORN motifs is sufficient for PM binding, but neither alone is sufficient for curvature sensing. To also determine the potential role of TM domain in curvature sensing, we generated a construct, JPH3-Sec61β, in which the TM domain of JPH3 was replaced with the transmembrane domain of Sec61β. Remarkably, JPH3-Sec61β exhibited similar curvature sensing capability compared to full-length JPH3 (**Fig. 5m, k, l**). This suggests that the TM domain itself does not influence the PM curvature preference of JPH3.

As neither the MORN motifs nor the LCR alone is sufficient to mediate curvature sensing, we constructed GFP-8MORN_LCR and GFP-8MORN_LCR-αHelix, which include 8 MORN motifs together with the joining LCR, with or without the αHelix. Interestingly, both GFP-8MORN_LCR and GFP-8MORN_LCR-αHelix exhibited PM localization and a clear preference for PM curvature at the nanobar ends (**Fig. 5n, o, k**). The distribution of 8MORN_LCR-αHelix is more uniform on the PM compared to GFP-8MORN_LCR, likely due to the stabilizing effect of the αHelix. Quantifications of the nanobar end-to-side ratios showed that the PM curvature preferences of 8MORN_LCR and 8MORN_LCR-αHelix are significantly higher than that of LCR and 8MORN-αHelix (**Fig. 5l**). Similarly, 8MORN_LCR-αHelix of both JPH2 and JPH4 exhibit PM binding and curvature targeting properties (**Fig. 5p, k, l**). Since Δ8MORN, ΔLCR, 8MORN-αHelix, and LCR show no curvature preferences, but 8MORN_LCR and 8MORN_LCR-αHelix both show strong curvature preference, we conclude that both the 8MORN motifs and the LCR are required for JPH’s curvature sensing.

Pathogenic variants in JPH2, such as S101R, Y141H, and S165F mutations, have been associated with hypertrophic cardiomyopathy^46^. Previous studies have shown that these mutants result in greatly impaired cellular calcium handling through unclear mechanisms^46^. Our measurements revealed that both S101R and S165F exhibit significantly impaired curvature localization, with reduced nanobar end-to-side ratios compared to wild-type (WT) JPH2 (**Extended Data Fig. 4b, c**). On the other hand, Y141H mutation did not affect the curvature sensing of JPH2 (**Extended Data Fig. 4b, c**). Our findings suggest that S101R and S165F, but not Y141H, may contribute to cardiac pathology through their reduced preference for PM curvature.

### 6. EHD proteins interact with JPHs and convey the PM-curvature preference

As JPHs do not harbor any known curvature-sensing domain, we hypothesized that JPHs’ curvature preference arises from its interactions with other PM proteins that are curvature sensitive. To identify such proteins, we analyzed the JPH2 interactome, previously obtained through proximity labeling techniques in a published study^47^, and selected those that are localized to the PM and are either intrinsically curvature-sensitive or participate in curvature-dependent processes such as endocytosis. Among more than 700 proteins in the JPH2 interactome, we identified seven proteins: clathrin heavy chain, EHD2, EHD4, and Caveolae-associated protein 1, 2, 3, and 4 (CAVIN1-4) as candidates (**Fig. 6a, Supplementary Table S1**). Of these proteins, previous research has established that EHDs and CAVINs are curvature-sensing proteins participating in caveolae formation ^48–52^ and T-tubule formation^53–56^; while clathrin is a crucial component of clathrin-mediated endocytosis. Clathrin preferentially accumulates at nanobar ends, as reported in a previous study ^22^. We further confirmed the PM curvature sensitivity of these proteins by showing that EHD4, CAVIN1, Caveolin1, and Caveolin2 all preferentially accumulated at the nanobar ends in U2OS cells (**Extended Data Fig. 5a**).

**Fig. 6:**
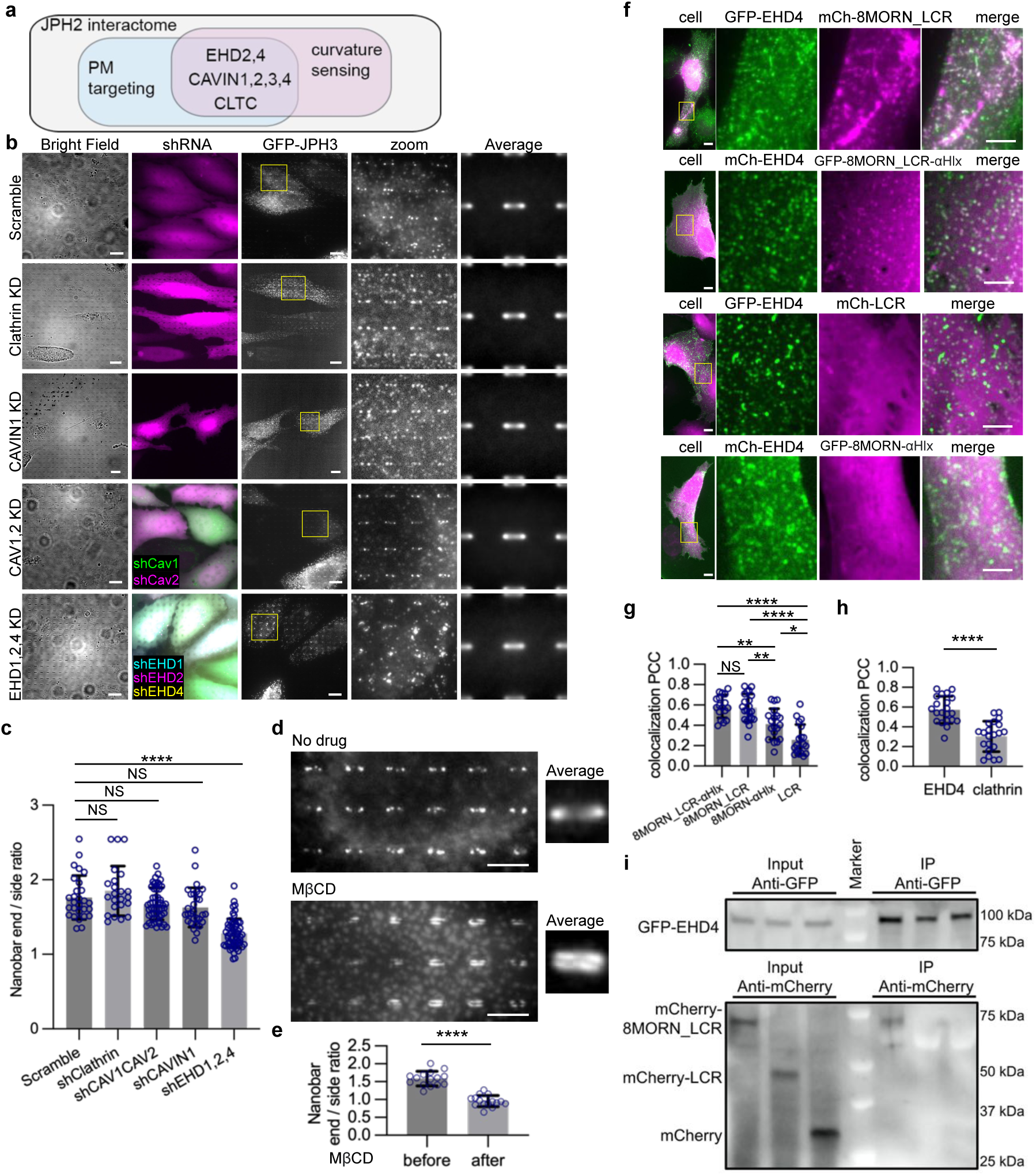
EHD proteins interact with JPHs and convey the PM curvature preference. **a**, An illustration of the overlapping between the PM targeting and the curvature sensing candidates in JPH2 interactome. **b**, Representative images of GFP-JPH3 on nanobars shRNA knockdown of scramble, Clathrin, CAVIN1, CAV1/2, or EHD1/2/4. ShRNAs sequences were fused with fluorescence proteins to identify knockdown cells. Bright field and the fluorescent protein expression are shown on the left. Zooms are the enlarged view of the region in yellow boxes. Averaged nanobar images from multiple cells are shown on the right. Cell number for average: scramble = 28, shCLTC (Clathrin heavy chain) = 23, shCAV1/2 = 53, shCAVIN1 = 30, shEHD1/2/4 = 55. Scale bar 10 µm in whole cell images, 5 µm in zoom-in images. **c**, Quantifications of GFP-JPH3 end-to-side ratios upon shRNA knockdowns. Conditions and cell numbers are the same as in b. ****P<0.0001, NS(shClathrin) P>0.9999, NS(shCAV1shCAV2) P>0.9999, NS(shCAVIN1) P=0.0592. **d**, The distribution of GFP-JPH3 on nanobars before and after 10 mM MβCD treatment at 37°C for 30 min. Scale bar 5 µm. Right: averaged images of all bars in a single cell. **e**, Quantifications of the nanobar end-to-side ratios for GFP-JPH3 in U2OS cells before and after the MβCD treatment. n = 15 cells for each group. **f**, Representative images of U2OS cells co-expressing EHD4-GFP/mCherry with indicated JPH3 mutants. Enlarged views are zoomed from the yellow boxes in whole cell image on the left. Scale bar 10 µm in whole cell images, 5 µm in zoom-in images. **g**, Pearson’s correlation coefficients (PCC) between coexpressed EHD4-GFP/mCherry and JPH3 mutants shown in (**f**). Cell number: mCherry-8MORN_LCR = 21, GFP-8MORN_LCR-αHelix = 17, mCherry-LCR = 19, GFP-8MORN-αHelix = 19. ****P < 0.0001, **P = 0.0024 (8MORN_LCR-αHlx vs. 8MORN-αHlx), **P = 0.0075 (8MORN_LCR vs. 8MORN-αHlx), *P = 0.0174 (8MORN-αHlx vs. LCR), NS P= 0.9995 (8MORN_LCR-αHlx vs. 8MORN_LCR). **h**, PCC for colocalization between mCherry-8MORN_LCR and EHD4-GFP or between mCherry-8MORN_LCR and anti-clathrin. n = 21 cells for each group. ****P<0.0001. **i**, Co-immunoprecipitation (IP) from HEK cells expressing EHD4-GFP and mCherry-8MORN_LCR or mCherry-LCR. GFP was pulled via GFP-antibody beads, and the immunoprecipitates were blotted with mCherry antibody. mCherry-8MORN_LCR, but not mCherry-LCR, was pulled together with EHD4-GFP. All experiments were independently replicated at least three times. All error bars represent STD. Kruskal–Wallis test corrected with Dunn’s multiple-comparison test was used for (**c**). Unpaired two-tailed Welch’s t test was used to assess significance for (**e**,**h**). Brown-Forsythe and Welch ANOVA tests was used to assess significance for (**g**).

To determine which proteins are crucial for JPH curvature sensing, we used small hairpin interference RNA (shRNA) to separately knock down clathrin heavy chain (gene CLTC), EHD-1/2/4, CAVIN1, or Caveolin-1/2 (gene CAV1, CAV2) isoforms that are expressed in U2OS cells (**Extended Data Fig. 5b**). Although caveolins were not detected in the JPH2 interactome, we included caveolins in our studies as they directly interact with and functionally coordinate with EHDs and CAVINs, and previous immunoprecipitation studies suggested a molecular interaction between JPH-1/2 and Caveolin-1/3 ^57–59^. Successful knockdown of these proteins was confirmed by western blotting (**Extended Data Fig. 5c**). We found that shRNA knockdown of clathrin heavy chain or CAVIN1, or double knockdown of Caveolin-1 and -2 did not affect the preferential accumulation of JPH3 at the nanobar ends (**Fig. 6b**) or their end-to-side ratios (**Fig. 6c**). On the other hand, we found that triple knockdown of EHD1, 2, and 4 significantly inhibited JPH3’s accumulation at nanobar ends (**Fig. 6b**) and reduced its end-to-side ratio (**Fig. 6c**). These knockdown studies indicate that the EHD family of curvature-sensing proteins is crucial for JPH3’s curvature preference.

When co-expressed, we found that different EHD isoforms, such as EHD1 and EHD4, or EHD2 and EHD4, colocalize extensively (**Extended Data Fig. 5d**). Because both EHD-2 and -4 are identified in JPH’s interactome and EHD4 shows a higher peptide count ^47^, we used EHD4 as a representative of the EHD family in our subsequent investigation. When co-expressed, EHD4-mCh and GFP-JPH3 both display clear preferential accumulation at nanobar ends (**Extended Data Fig. 5e**), indicating that EHD4 is located at JPH3-mediated contact sites at curved PMs. However, it is necessary to point out that EHD4 and JPH3 only partially co-localize, likely due to their involvement in other independent cellular processes.

In addition to U2OS cells, we also confirmed the curvature preference of EHDs and their functional roles in ER-PM contacts in cardiomyocytes. Immunofluorescence staining of EHD2 and EHD4 in both iPSC-CMs and rat embryonic CMs showed significant enrichment at curved PM regions surrounding nanopillars (**Extended Data Fig. 5f**). Quantification of the nanobar end-to-side ratio of EHD4 in iPSC-CMs further confirmed the significant curvature preference of EHDs (**Extended Data Fig. 5g**). Similar as in U2OS cells, triple knockdown of EHD1, EHD2, and EHD4 in iPSC-CMs resulted in a significant reduction of the end-to-side ratio of JPH2 (**Extended Data Fig. 5h**).

The proper localization of EHD protein can be significantly disrupted by cholesterol extraction using methyl-β-cyclodextrin (MβCD) ^60^. On nanobars, EHD4-GFP exhibited a drastic loss of PM curvature preference after MβCD treatment (**Extended Data Fig. 6a**). Upon cholesterol depletion by MβCD, we also observed a profound relocalization of GFP-JPH3 from the nanobar ends to the sides (**Fig. 6d**). The averaged nanobar image transitioned from a dumbbell shape to two parallel lines alongside the walls of the nanobars, and the nanobar end-to-side ratio decreased from 1.6 to 0.95 (**Fig. 6e**). These results further support the role of EHD proteins in JPH3’s PM curvature preference.

Given the clear preference of the 8MORN_LCR domain for PM curvature, we hypothesized that EHDs may interact with this specific region of JPH3. To investigate this hypothesis, we separately co-expressed 8MORN_LCR, 8MORN_LCR-αHelix, 8MORN-αHelix, and LCR with EHD4 in U2OS cells. We observed that EHD4 formed puncta or tubular structures on the PM (**Fig. 6f**), which are typical of curvature-sensing proteins and agree with previous studies^48,49^. Interestingly, both 8MORN_LCR and 8MORN_LCR-αHelix strongly colocalize with EHD4 in these punctate structures but 8MORN-αHelix and LCR appear to uniformly distributed (**Fig. 6f**). In EHD1/2/4 triple knockdown cells, ΔLCR still form ER-PM contacts similar to control cells, confirming that the MORN domains bind to the PM independent of EHDs (**Extended Data Fig. 6b**). Quantification of the Pearson Correlation Coefficients revealed a significantly stronger correlation between EHD4 and 8MORN_LCR or 8MORN_LCR-αHelix compared to between EHD4 and 8MORN-αHelix or LCR. These observations indicate that both 8MORN and LCR are necessary for interaction with EHD4. It is worth noting that 8MORN-αHelix displays stronger colocalization with EHD4 than LCR (**Fig. 6g**), despite having weaker PM affinity. This suggests a more important role of MORN motifs in EHD interaction. As a control, mCherry-8MORN_LCR did not colocalize with clathrin-marked puncta structures on the PM (**Fig. 6h, Extended Data Fig. 6c**).

To further determine whether EHD4 interacts with 8MORN_LCR, we carried out co-immunoprecipitation studies. GFP-EHD4 was co-expressed with either mCherry-8MORN_LCR, mCherry-LCR, or mCherry, in HEK293T cells. When GFP-EHD4 was pulled-down with anti-GFP-antibody-conjugated beads, we found that mCherry-8MORN_LCR co-precipitated together with GFP-EHD4, but mCherry-LCR or mCherry did not (**Fig. 6i**). Overall, our results support a molecular model in which the curvature preference of JPH-mediated ER-PM contact is due to interactions between the N-terminal segment of JPH3 and EHDs.

## Discussion

ER-PM contacts were first discovered in the 1950s in striated muscle cells that have extensive T-tubule systems^61^. The ER-PM contacts are a prominent feature of striated muscle cells, as they occupy up to 50% of the T-tubule membranes^17^. In comparison, ER-PM contacts only occupy 4-8% of the external plasma membrane and even less in non-muscle cells. Surprisingly little is known about the mechanism of such enrichment of ER-PM contacts on T-tubule membranes. Our results suggest a mechanism by which cells can harness the PM curvature of T-tubules to spatially enrich ER-PM contacts. By using vertical nanobars and showing that ER-PM contacts preferentially form at the nanobar ends compared to the sidewalls, we conclusively demonstrate that T-tubules enrich ER-PM contacts by the nature of their PM curvature, not by simply bringing the PM closer to the ER.

Our observation that PM curvature enriches JPH-mediated ER-PM contacts provides a new perspective on the role of membrane curvature in ER-PM contact formation. Recent studies have shown that ER membrane curvature might play a role in lipid transfer at ER-PM contacts in yeast ^62,63^. The asymmetric packing of lipid molecules in the outer leaflet of the curved membrane is long believed to accelerate lipid exchange ^64^. Our finding that PM curvature promotes local ER-PM contact formation through curvature-sensing proteins may inspire new discoveries about their effect on lipid transfer functions aside from Ca^2+^ signaling.

Our study reveals that a polybasic sequence in the JPH’s LCR region has a high affinity for the PM, while the MORN motifs have a lower affinity for the PM. When LCR was expressed without the ER-transmembrane domain, they were strongly localized to the PM, likely through electrostatic interactions with negatively phospholipids on the PM. Mutations of the polybasic residues completely abolished their PM localizations. The electrostatic interaction may, at least partially, explain recent observations that show JPH2 nuclear localization upon calpain-2-mediated C-terminal cleavage^65–67^.

Importantly, our studies reveal that the MORN_LCR motifs interact with curvature-sensing EHDs on the PM, which can recruit JPHs at PM curvatures. The disruption of EHD localization by MβCD cholesterol extraction disturbed the curvature preference but not the PM targeting of JPH3. This suggests two populations of JPH3-mediated ER-PM contacts: one positive with EHDs/cholesterol and the other independent from EHDs/cholesterol, which is in agreement with the previous observation that JPHs cofractionate with both the caveolin-rich lipid rafts and the non-lipid-raft domains^68^.

In our studies, PM curvatures were generated by vertical nanostructures, whereas in vivo, PM curvatures are generated and stabilized by proteins. For example, Bridging Integrator 1 (BIN1), a curvature sculpturing and sensing protein, is known to be essential for T-tubule generation and stabilization^69,70^. We found that BIN1 knockdown by shRNAs (**Extended Data Fig. 5c**) slightly reduced the curvature preference of JPHs, although the effect was not as significant as that of EHD knockdowns (**Extended Data Fig. 7**). The potential coordination and synergy between EHDs and BIN1 in this process warrants further investigation.

Previous research has demonstrated that EHD proteins generate membrane tubules with a diameter range from 20 to 100 nm^48,49^. In contrast, T-tubules of cardiomyocytes have a mean diameter of ∼250 nm in rodents and ∼ 400 nm in larger mammals^71^, surpassing the curvatures typically generated by EHDs. However, in our studies, EHDs show a clear curvature preference to nanopillars and nanobars, which induce PM curvatures with a diameter range of 200-400 nm (**Extended Data Fig. 5a, 6a**). It is worth noting that while curvature sensing and curvature generation are often properties of the same proteins^72^, they may involve different molecular interactions - with curvature sensing dominated by protein–membrane interactions, while curvature generation involving protein oligomerization and scaffolding in addition to protein-membrane interaction^73–75^. Therefore it is plausible that the same proteins may generate and sense membrane curvatures at different diameter ranges. For example, another curvature-sensing protein, FBP17, induces membrane tubules with an average diameter of ∼ 70 nm but can sense membrane curvatures with a diameter up to 450 nm^76,77^. Nevertheless, our study is based on observations of nanostructures of 200-300 nm in diameter. The curvature preferences of ER-PM contact proteins like JPHs and E-Syt2s beyond this curvature range are yet to be explored.

JPHs and E-Syts are both ER-PM tethering proteins, but they show distinct preferences for PM curvatures. This may contribute to a previous observation in *C. elegans* that JPHs and E-Syts exhibit different localization in presynaptic sites in the dorsal cord and display antagonistic effects on synaptic transmission^78^. Moreover, while E-Syts are broadly expressed, JPHs are specifically expressed in excitable cells (muscle cells, neurons, T cells^79^, and pancreatic beta cells^80^), in which calcium dynamics are critical for their excitability. For example, JPH1 and JPH2 are expressed in muscle cells, while JPH3 and JPH4 are expressed in neurons and are important for the generation of neuronal afterhyperpolarization currents^81,82^. In our studies, several JPH2 mutations associated with hypertrophic cardiomyopathy result in the impairment of JPH2’s targeting to membrane curvatures, which likely contributes to the defects in calcium handling and hypertrophic cardiomyopathy in these patients. Aside from the mutations we examined, dozens of mutants in JPH2 have been clinically associated with hypertrophic cardiomyopathy, dilated cardiomyopathy, arrhythmias, and sudden cardiac death^7,83,84^. Furthermore, toxic aberrant transcription and loss of expression of JPH3 have been implicated to play a role in Huntington’s disease-like 2 (HDL2) pathophysiology^85^. Our findings on the molecular mechanisms of JPHs will shed light on related mechanistic and therapeutic studies.

The PM curvatures have been shown to affect a range of PM processes from clathrin-mediated endocytosis, integrin adhesions, glycoprotein and ion channel distributions to actin dynamics. Our finding that PM curvature directly regulates ER-PM contacts and thus cellular calcium responses opens a new frontier in this exciting area.

## Methods

### Nanopillar and Nanobar Fabrication

In the sample preparation process, fused quartz substrates were meticulously cleaned using acetone and isopropanol, followed by sonication to eliminate contaminants. After thorough cleaning, the coverslips were dried at 180°C. Subsequently, a 275 nm layer of 9% CSAR 62 E-beam resist was spin-coated onto the substrate and baked at 180°C for 3 minutes, and a 100 nm conductive Electra 92 layer was applied and baked for an additional 3 minutes. Using the Raith Voyager lithography system, nanopatterned pillars and BAR with varying dimensions were fabricated while compensating for proximity effects. After lithography exposure, the Electra 92 conductive layer was removed in deionized water, and xylene development revealed the nanopatterns. A 120 nm chromium (Cr) masking layer was evaporated onto the substrate, followed by plasma etching using the PlasmaTherm Versaline LL ICP Dielectric Etcher to reach a depth of 1500nm. Last, the Cr layer was removed with a Cr etchant, resulting in the desired nanopatterns. The substrate and its dimensions were characterized using Magellan Scanning Electron Microscopy (SEM).

### Cell Culture

U2OS cells (ATCC HTB-96™), HeLa cells (ATCC CCL-2™) and HEK293T cells (ATCC® CRL-3216™) were maintained in DMEM (Gibco 11965-092) supplemented with 10% (v/v) fetal bovine serum (FBS) (Sigma F4135). Cells were passaged every 3-5 days to maintain sub-confluency. iCells (Fujifilm Cellular Dynamics, 01434), dissected rat embryonic CMs, and iPSC-CMs were maintained in RPMI (Gibco REF 11875-093) with B-27 supplement (50:1, Gibco REF 17504-044). HL-1 cells were obtained from the laboratory of William C. Claycomb at Louisiana State University, and maintained in Claycomb medium supplemented with 10% FBS. Medium for iPSC-CMs, iCells, rat embryonic CMs and HL-1 cells were changed every 2-3 days.

### Induced Pluripotent Stem Cell Derived Cardiomyocyte (iPSC-CM) Differentiation

Human induced pluripotent stem cells (iPSCs) were obtained from Stanford Cardiovascular Institute and cultured in E8 medium (Gibco, Life Technologies) on 6-well plates coated with Matrigel (Corning). Cells were passaged at a 1:12 ratio after 5 min of incubation with Accutase (Sigma-Aldrich) at 37°C. After replating, the E8 medium was supplemented with 10 μM Y-27632 ROCK inhibitor (Selleck Chemicals) for 24 hours. Subsequently, the medium was changed to standard E8 medium with daily medium changes. For differentiation into CMs, hiPSCs were seeded at a 1:12 ratio and cultured until they reached 85% confluence. Differentiation was initiated by changing the medium to RPMI supplemented with B27 without insulin (Life Technologies) and 6 μM CHIR-99021 (Selleck Chemicals). At 48 hours post-induction, the medium was switched to RPMI-B27 without insulin for 24 hours and then supplemented with 5 μM IWR-1 (Selleck Chemicals) for another 48 hours. Metabolic purification of CMs was conducted on day 11 using RPMI-B27 without D-glucose (Life Technologies) for 96 hours. After purification, CMs were maintained in RPMI-B27 for future experiments.

### Cell Culture on Nanochips

Before seeding cells on nanochips (both nanopillar chips and the nanobar chips were treated the same way), we treated the nanochips with air plasma (Harrick Plasma) at high power for 10 min, then coated the nanochips with 0.1 mg/ml poly-L-lysine (PLL) in phosphate-buffered saline (PBS) at room temperature (RT) for 30 min. The chips were then washed with PBS 3 times before incubation with 0.5% (v/v) glutaraldehyde (Sigma-Aldrich, G6257) in PBS at RT for 10-15 min, and followed by another 3 washes with PBS wash and incubation with the desired ECM protein. Specifically, for U2OS, HeLa, and HEK293T cells, the chips were incubated with 0.1% (m/v) gelatin combined with 20 µg/ml human plasma fibronectin (Sigma-Aldrich, 341635) in PBS at 37°C for 1 hour, then followed by 3 washes with PBS. If fluorescent imaging was needed later, the chips were then incubated with 5 mg/ml sodium borohydride in PBS at room temperature for 5 min to quench the autofluorescence from glutaraldehyde followed by 3 washes with PBS before seeding cells. For iPSC-CMs, rat embryonic CMs, and iCells, the ECM protein used was Matrigel(Corning REF 356231). Matrigel was diluted with ice cold DMEM/F12 + GlutaMAX (Gibco REF 10565-018) at a 1:200 ratio, and added to the chips to incubate at 37°C for 1 hour overnight. Cells were then seeded on chips after removing the extra Matrigel. The seeding medium used for iPSC-CM was RPMI+B-27+10% KnockOut SR (KSR) Multi-Species (Gibco REF A31815-01). The medium was changed to maintenance medium after 24 hours.

### Rat Embryonic CM isolation

Use of laboratory animals approved by the Stanford University administrative panel on laboratory animal care, and complies with all of the relevant ethical regulations.

Freshly dissected hearts from E18 rats were cut into 4 pieces, and then washed with HHBSS buffer (HBSS (Gibco, 14025126) +10 mM HEPES +1 mM glucose). Cardiomyocytes were isolated in Triple Select 10x (Gibco REF A12177-01) for 30 min at 37 ℃ with agitation. Isolated cardiomyocytes were cultured in 10% KSR in RPMI supplemented with B27 overnight. The medium was changed to RPMI+B27 maintenance medium afterwards.

### Antibodies and Reagents

Anti-Junctophilin2 antibody (HPA052646), anti-α-actinin antibody (A7811), anti-Cav1.2 antibody (C1103), and anti-BIN1 antibody (SAB1408547) were purchased from Sigma. Anti-RyR2 antibody (NB1202827) was purchased from Novus Bio. Anti-Caveolin1 antibody (sc-70516) was purchased from Santa Cruz. Anti-Caveolin2 antibody (CSB-PA004572LA01HU-20UG), anti-EHD1 antibody (CSB-PA884470LA01HU-20UG), and anti-EHD2 antibody (CSB-PA873710LA01HU-20UG) were purchased from Cusabio. Anti-EHD4 antibody (50-172-7111) was purchased from Fisher Scientific. Anti-Clathrin antibody (MA1-065), anti-Calreticulin antibody (PA3-900), and anti-GFP antibody (A-11122) were purchased from Invitrogen. HRP-linked goat anti-rabbit IgG (H+L) antibody (7074) and HRP-linked goat anti-mouse IgG (H+L) antibody (7076) were purchased from Cell Signaling Technology. Anti-mCherry antibody (SAB2702291), methyl-β-cyclodextrin (C4555), and thapsigargin (Tg) (T9033) were purchased from Sigma.

### Plasmids

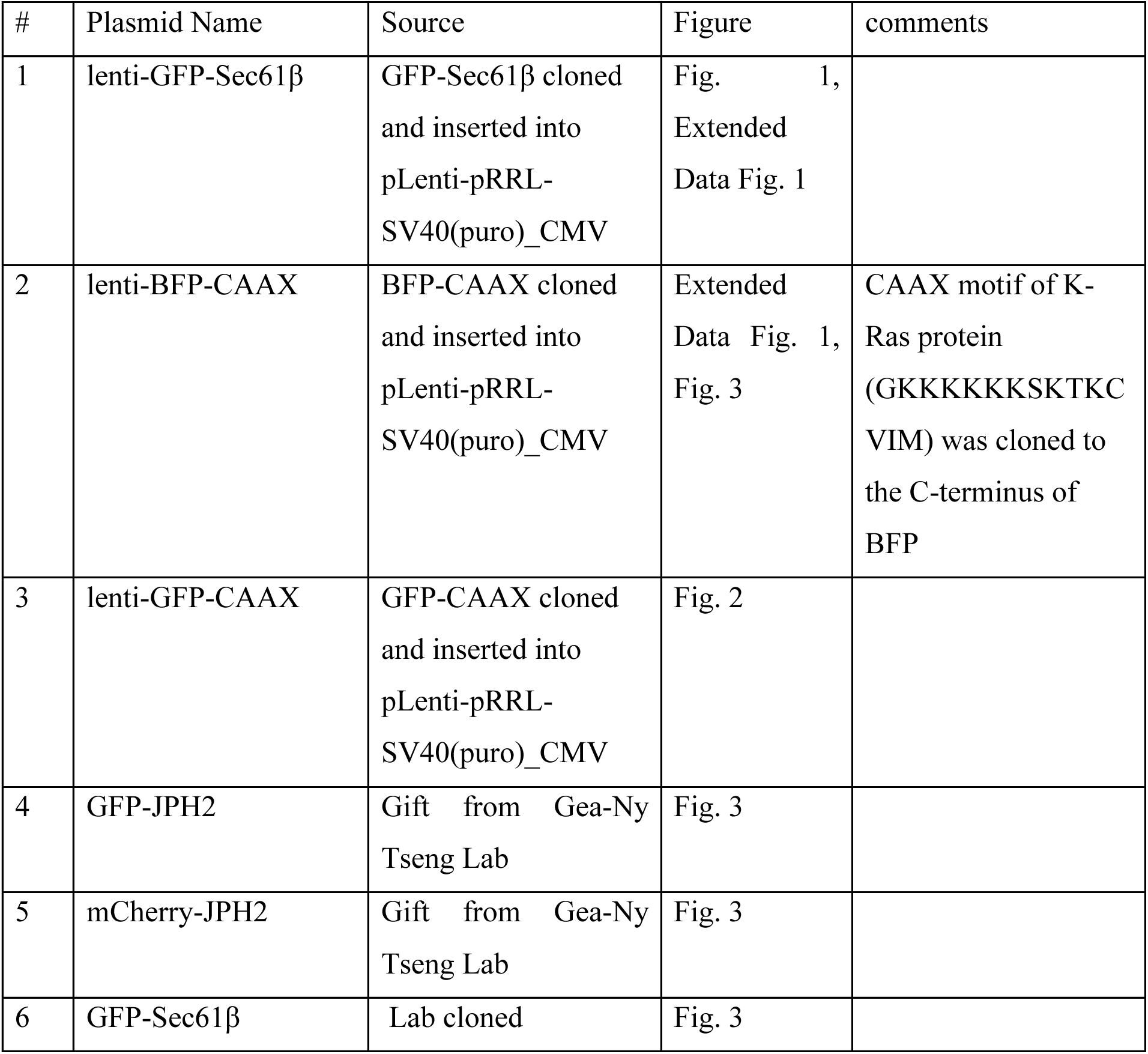
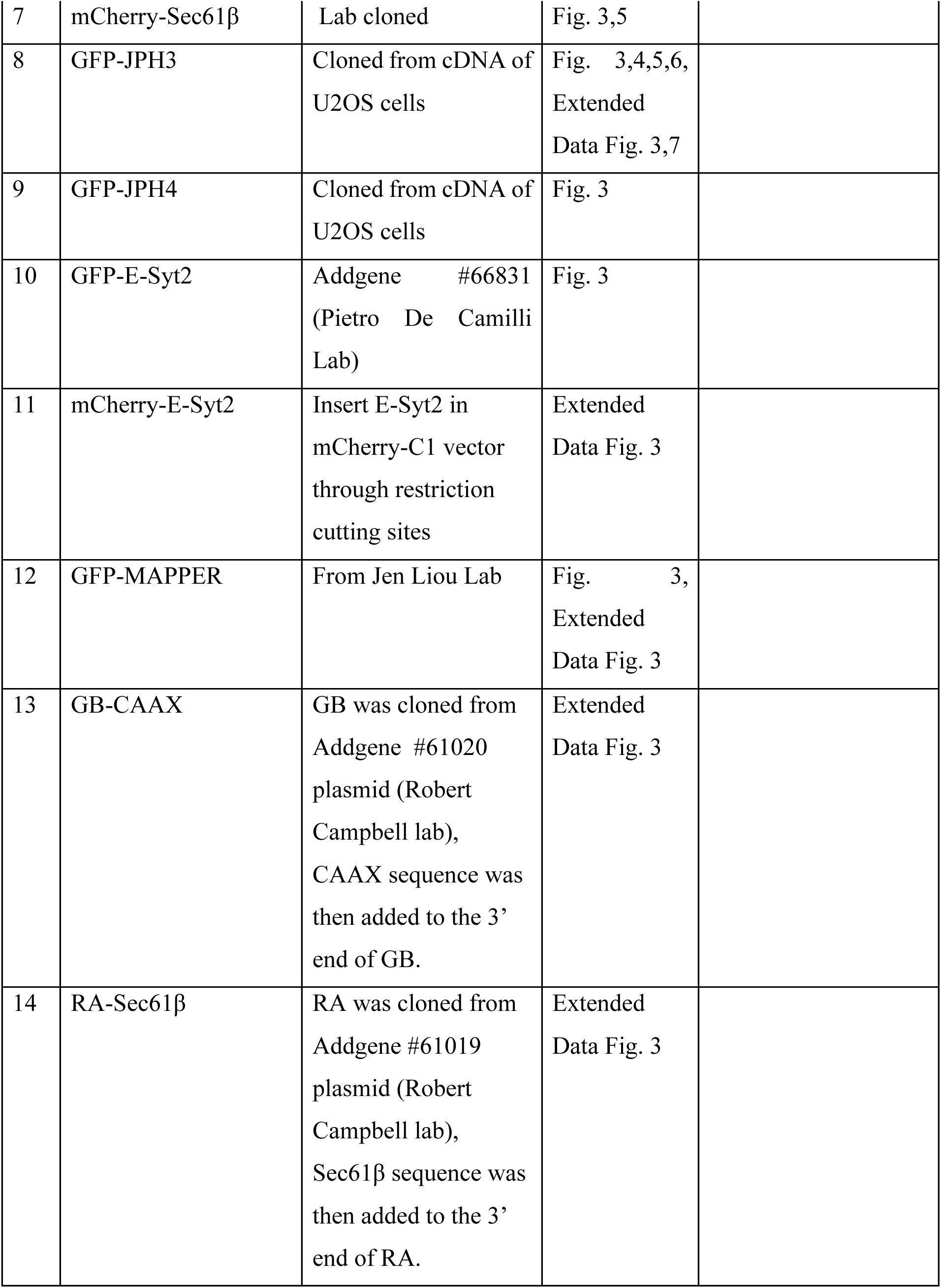
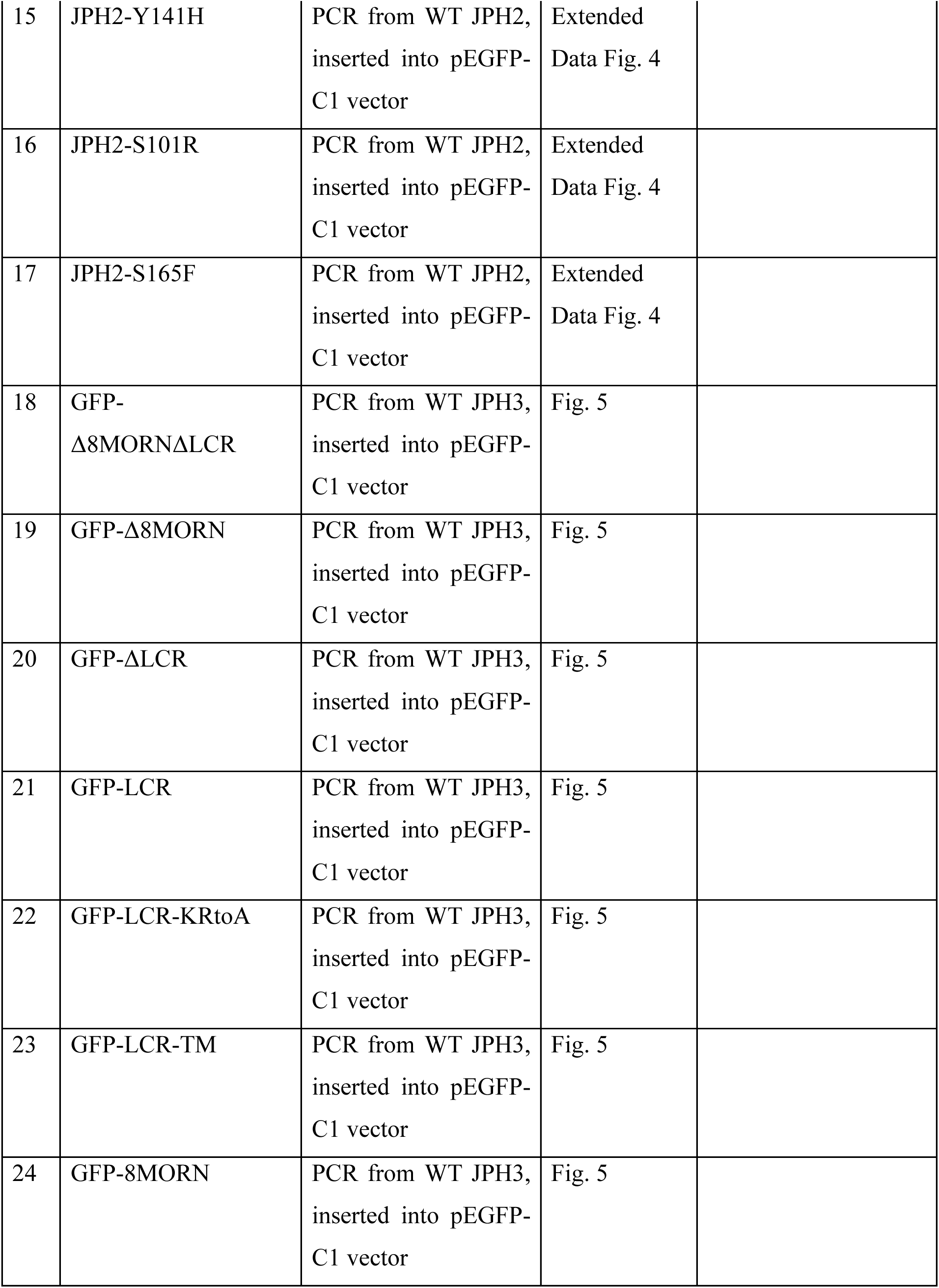
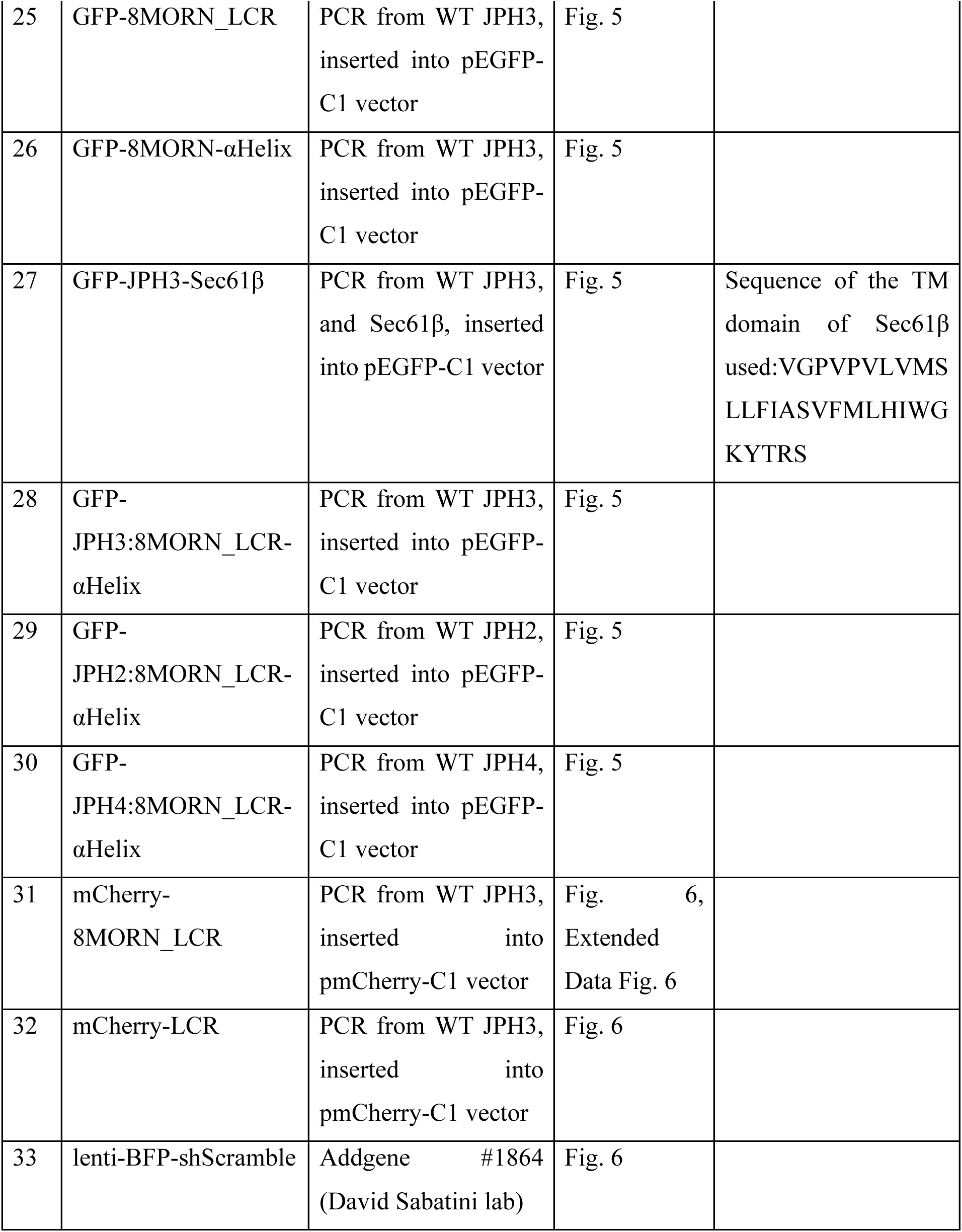
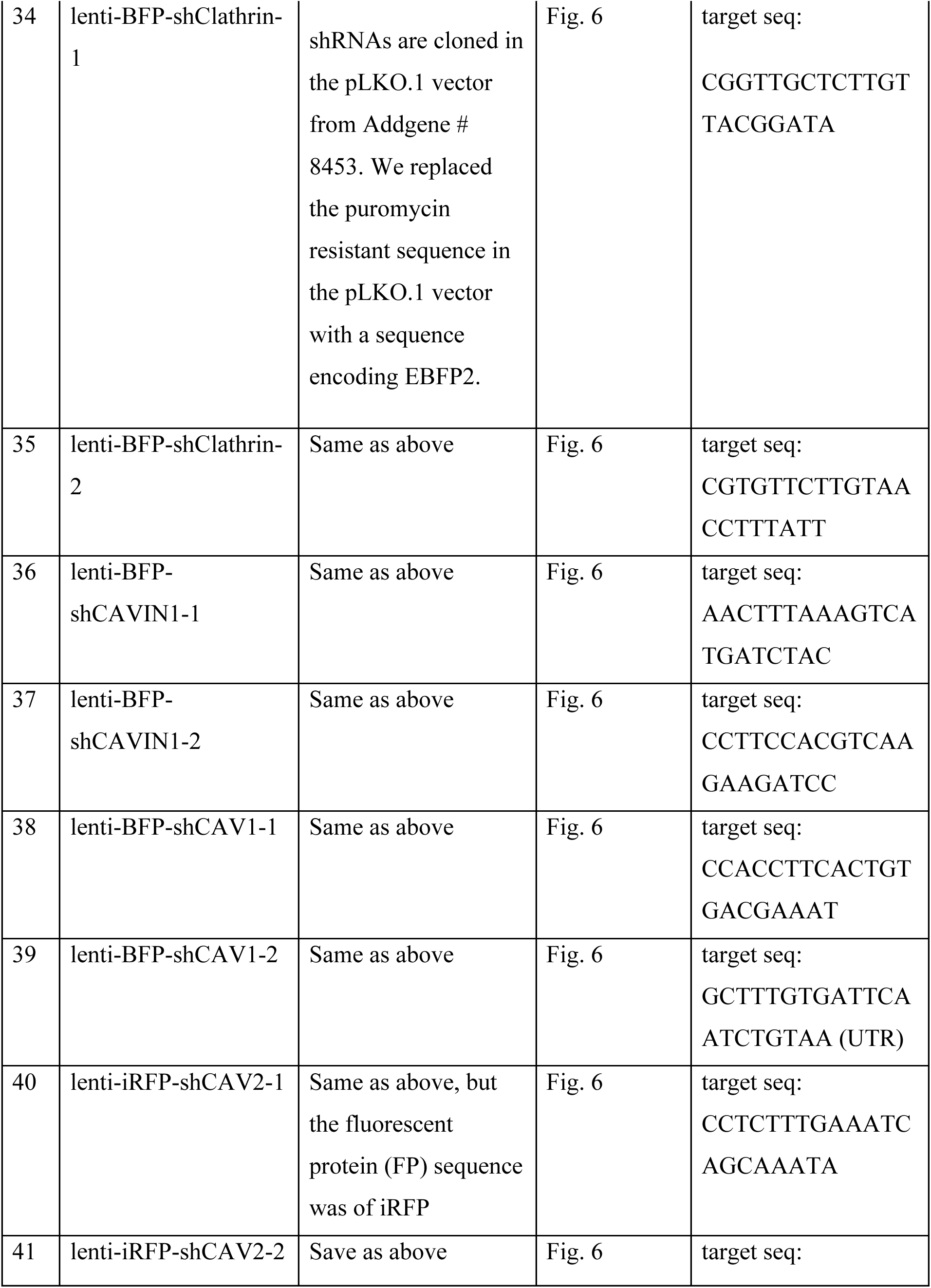
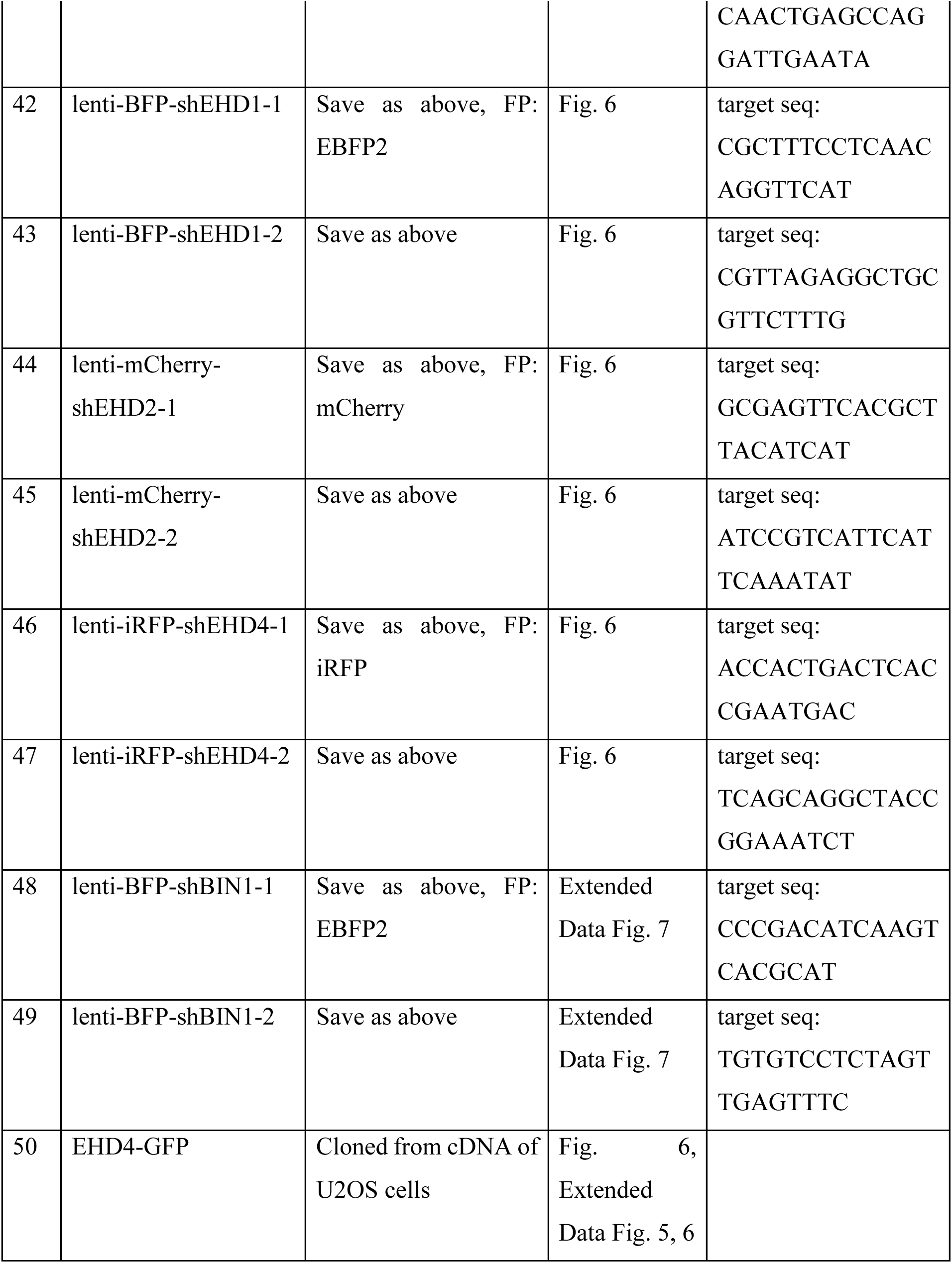
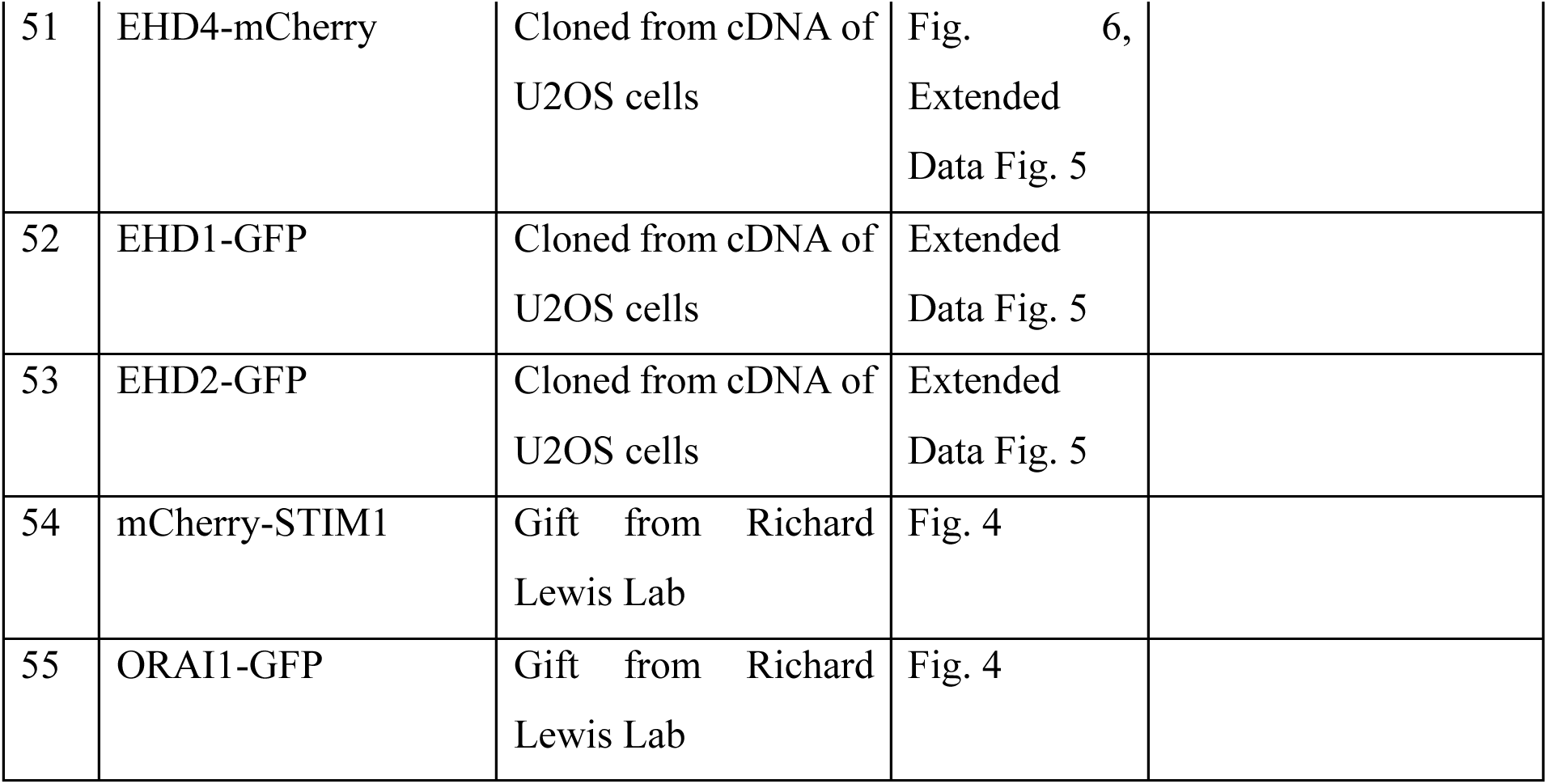

The plasmids we made in this research are all readily available upon request to the corresponding author.

### Cell Transfection

For imaging experiments, U2OS cells and HEK293T cells were transfected through electroporation (Lonza Amaxa Biosystems Nucleofector II) using pre-installed protocols. For each transfection, ∼0.5 million cells were electroporated with 0.2-0.5 µg plasmid DNA in 100 µL Electroporation buffer II (88 mM KH_2_PO_4_ and 14 mM NaHCO_3_, pH 7.4) freshly supplemented with 2 µL Electroporation buffer I (360 mM ATP + 600 mM MgCl_2_). HeLa cells were transfected with Lipofectamine 2000 (Invitrogen 11668-019). Plasmid DNA (0.5-1 µg) was used for each transfection of 0.2 million cells according to the reagent protocol.

### Lentiviral Particle Packaging

Lentivirus was generated in HEK293T cells plated at ∼70%-80% confluency in 6 well plates. Prior to transfection, the medium was switched from DMEM+10% FBS to pre-warmed DMEM. Then, 0.8 µg psPAX2 plasmids(Addgene #12260), 0.7 µg pMD2.G plasmids (Addgene #12259), and 1.5 µg of transfer plasmid encoding targeted cDNA or shRNA were transfected into 1 well of cells using lipofectamine 2000. The transfection media was switched to DMEM+10%FBS+1 mM Sodium Pyruvate (Gibco 11360). The transfected cells were cultured for 24 h before the virus-containing medium was collected, filtered through 0.45 µm PVDF filters (Millipore) and added to targeted cells. Lentiviral particles containing cDNA of GPF-CAAX were used to infect iPSC-CMs, and lentiviral particles containing shRNAs were used to infect U2OS cells to induce protein knockdown.

### shRNA Interference Experiments

shRNA coding sequences were cloned and inserted into 3rd generation transfer plasmid pLKO.1-TRC cloning vector (Addgene #10878) following the Addgene protocol. The sequences of shRNA oligos were either from Sigma-Aldrich’s predesigned shRNA or from Addgene as listed in the plasmid table. Each shRNA was cloned and inserted into a vector expressing a fluorescent protein to monitor transfection efficiency. The knockdown effects were examined on day 3 after lentiviral transfection of shRNAs. In the case of double or triple knockdown (KD) cells, fluorescent proteins of different colors were employed.

### Immunoblotting

U2OS cels 4 days after shRNA-lentiviral infection were lysed in RIPA buffer (25 mM Tris–HCl, 150 mM NaCl, 1% Triton X-100, 1% sodium deoxycholate, 0.1% sodium dodecyl sulfate) supplemented with protease and phosphatase inhibitor cocktails (Roche, 04693159001 and 04906837001) for 30 min on ice with vortex agitation every 5-10 min. The lysate was then centrifuged at 12000 rpm at 4°C for 10 min. The supernatants were then boiled at 95°C for 10 min and subjected to sodium dodecyl sulfate–polyacrylamide gel electrophoresis (SDS-PAGE). After electrophoresis, the samples were transferred onto nitrocellulose membranes using the Trans-Blot Turbo Transfer System (Bio-Rad, 1704150). The membranes were stained with Ponceau solution (5% v/v glacial acetic acid and 0.1% w/v Ponceau S) to confirm the equal loading then blocked with 5% milk in TBS-T buffer and then incubated with the indicated antibody diluted in 3% BSA in TBS-T at 4°C overnight. The protein bands were visualized using HRP-conjugated secondary antibody and chemiluminescence under Azure Imaging Systems (Azure Biosystem).

### Co-Immunoprecipitation (Co-IP) Assay

HEK293T cells (ATCC® CRL-3216™) were co-transfected with mCherry-8MORN_LCR, mCherry-LCR or mCherry alone together with GFP-EHD4 via electroporation and then incubated at 37°C overnight. Confluent cells were trypsinized and then harvested by spinning. After one wash with ice-cold 1X PBS, the cells were mixed with freshly prepared 0.5 mM DSP (3,3’-dithiobis (succinimidyl propionate), Sigma-Aldrich) in 1X PBS for 40 min at room temperature. To quench the reaction, an ice-cold 25 mM Tris-HCl buffer was then added to the cell mixtures for 20 min. After centrifugation, cell pellets were lysed in 1X RIPA buffer supplemented with protease (cOmplete, Roche) and phosphatase inhibitors (PhosSTOP, Roche) for 1 hr at 4°C. The lysates were then subjected to sonication at a pulse of 20-25% amplitude for 10 s followed by a 25 s incubation on ice. The sonication–incubation cycle was repeated twice. Subsequently, the lysates were mixed with equilibrated GFP-Trap magnetic agarose (ChromoTek) and incubated overnight at 4°C. The bead-protein complexes were harvested by centrifugation and washed with 1X PBS for 5 times. To elute and denature proteins, the bead-protein pellets were subsequently resuspended in a 2X Laemmli sample buffer (with β-mercaptoethanol) and boiled for 5-10 min at 95°C. The samples were subjected to SDS-PAGE and Western blotting. The blots were incubated with anti-GFP and anti-mCherry in 1X TBS-T buffer supplemented with 3% BSA overnight at 4°C. The protein signals were visualized using HRP-conjugated secondary antibodies and chemiluminescence.

### Immunofluorescence

iPSC-CMs were seeded on nanochips coated with Matrigel as described in the previous sections. Each nanochip had both nanopillar regions and flat regions on the same chip, ensuring imaging of both flat surfaces and nanopillar surfaces under the same conditions. Cells were allowed to grow on nanochips for 3-4 days. For experiments in which GFP-CAAX was transfected into iPSC-CMs, the cells were allowed to grow on nanochips for 1 day in seeding medium and then the medium was switched to GFP-CAAX lentivirus harvesting medium. After 24 hours, the medium was changed to iPSC-CM maintenance medium and incubated for another 24 hours for more GFP-CAAX expression. The samples were then fixed with 4% paraformaldehyde for 15 min at RT, incubated with 5 mg/ml sodium borohydride in PBS at RT for 5 min to quench the autofluorescence, permeabilized with 0.1% Triton X-100 in PBS for 10 min at RT, and blocked with 3% BSA in PBS for 30 min at RT. Three PBS washes were performed in between each step. The samples were then incubated with the indicated primary antibodies against JPH2 (1:300), RyR2 (1:500), Cav1.2 (1:300) or α-actinin (1:1000) diluted in 3% BSA at 4°C overnight. The samples were then washed with PBS for 5 min for 3 times and then stained with the secondary antibodies for 1 hour at RT. Alexa Fluor^TM^ 594-goat anti-rabbit IgG (Invitrogen A11012) and Alexa Fluor^TM^647-goat anti-mouse IgG (Invitrogen A32728) secondary antibodies were used at a 1:1000 dilution in 3% BSA in PBS. Samples were subjected to fluorescence imaging or expansion microscope processing afterwards.

### Focused Ion Beam Scanning Electron Microscopy (FIB-SEM) Imaging

The procedure was adapted from a prior study by our group^29^. Substrates containing cells were first rinsed with 0.1 M sodium cacodylate buffer (Electron Microscopy Sciences) and fixed at 4 °C overnight using 3.2% glutaraldehyde (Sigma Aldrich). Subsequently, specimens underwent post-fixation with 4% osmium tetroxide and 2% potassium ferrocyanide (Electron Microscopy Sciences) for 1 h, followed by 1% thiocarbohydrazide (Electron Microscopy Sciences) for 20 min and 2% aqueous osmium tetroxide for 30 min. After rinsing twice with distilled water, samples were incubated overnight with syringe-filtered 4% aqueous uranyl acetate (Electron Microscopy Sciences, en-bloc step). Dehydration was achieved through an increasing ethanol series (10%– 30%–50%–70%–90%–100%, each step lasting 5–10 min), followed by specimen infiltration with epoxy-based resin using various ethanol-to-resin ratios (1:3 for 3 h, 1:2 for 3 h, 1:1 overnight, 2:1 for 3 h, and 3:1 for 3 h). Finally, samples were infiltrated with 100% resin overnight at room temperature, and after removal of excess resin, polymerization was carried out at 60 °C overnight.

Prepared samples were metal-sputtered and loaded in a dual-beam Helios Nanolab 600i FIB-SEM (FEI) vacuum chamber. Secondary SEM imaging used 3–5 kV voltage and 21 pA–1.4 nA current. Cross-section imaging employed a beam acceleration with 2–10 kV in voltage and 0.17–1.4 nA in current using a backscattered electron detector. Preservation of regions of interest involved double platinum layer deposition via in situ sputtering. Trenches were etched at 30 kV acceleration voltage and 9.1–0.74 nA current followed by a fine polishing of the resulting cross-sections using 30 kV voltage and 80 pA current.

### Expansion Microscopy (ExM) Imaging

Expansion microscopy was performed as previously described^31^. Cells were cultured on nanopillar chips, fixed and immunostained as described above. The samples were then incubated overnight at RT in a 1:100 dilution of AcX (Acryloyl-X, SE, 6-((acryloyl)amino) hexanoic acid succinimidyl ester, Invitrogen A20770) in PBS. The samples were then washed with PBS followed by a 15 min incubation at RT in gelation solution (19% (w/w) sodium acrylate, 10% (w/w) acrylamide, 0.1% (w/w) N,N’-methylenebisacrylamide). Gelation solution supplemented with 0.5% N,N,N’,N’-tetramethylenediamine (TEMED) and 0.5% ammonium persulfate (APS) was then prepared and briefly vortexed. The gelation solution, TEMED and APS were kept on ice before combining and APS was added last to initiate gelation. Nanochips were then flipped cell side down onto a 70 μL drop of this supplemented gelation solution on parafilm and incubated at 37°C for 1 h. The nanochip with the hydrogel still attached was then incubated in a 1:100 dilution of proteinase K in digestion buffer (50 mM Tris HCl (pH 8), 1 mM EDTA, 0.5% Triton X-100, 1 M NaCl) for 7 h at 37°C. The hydrogel at this point has expanded and detached from the nanochip. The hydrogel was then soaked twice in Milli-Q water for 30 min and then left to incubate overnight in Milli-Q water at 4°C. Before imaging, excess water on the hydrogels was carefully removed using a Kimwipe. The hydrogels were then mounted onto PLL-coated glass coverslips cell side down to prevent sliding during imaging.

### ExM Analysis

The expanded sample was imaged with a 40x water lens under a confocal microscope. Z-stack images were taken every 0.3 µm with 1.2 Airy Unit. The pixel size in the final image was 0.0777 in both x and y and 0.3 µm in z. The z direction was scaled up ∼3.86 (=0.3/0.0777) times for display (Fig.2d,f,g) so that the x and z were visually at the same dimensional scale. The x-z view displayed in Fig.2f was the average projection of the x-z view of 5 neighboring pixels in x, while each single pillar spanned approximately 20 pixels under an expansion microscope.

To quantify the pillar-flat ratio (Fig.2h), background was subtracted for both CAAX and JPH2 channels and single pillars were identified using the in-house MATLAB code for nanopillar analysis. To quantify intensities at each pillar and the flat region near each pillar, images were average-projected in x-z directions at each identified pillar. Specifically, for pillar intensity quantification, the average intensity of the 25 consecutive slices centered on each pillar’s central axis was quantified as the pillar intensity of that pillar, and 5-6 pillars in each row were averaged as one data point. For flat intensity quantification, the average intensity of 40 consecutive slices of the flat region near each pillar was quantified, and 5-6 such regions in each row were averaged as one data point. To obtain the ratio displayed in Fig. 2h, the ratio between the pillar intensity data points and the flat intensity data points at each pillar was first quantified. This ratio was larger than 1 even for CAAX due to the larger illuminating volume in z in the confocal point spread function. To normalize this effect, all the ratios from the last step were normalized by the average value of the ratio of the CAAX channel so that the averaged pillar-flat ratio of CAAX was one. The values after normalization are displayed in Fig. 2h.

To quantify how many fold JPH2 density was increased from the flat surface to the pillar surface, we quantified the ratio of JPH2/CAAX at the pillar over the JPH2/CAAX at the flat surfaces for each data point. The resulting fold change was 3.8 ±1.4 as reported in the main text.

### Fluorescence Imaging

Transfected U2OS cells, HeLa cells and HEK293T cells were imaged live in Ringer’s buffer (155 mM NaCl, 4.5 mM KCl, 2 mM CaCl_2_, 1 mM MgCl_2_, 5 mM HEPES, 10 mM D-glucose, pH=7.4), and the fixed cells were imaged in PBS at RT unless otherwise stated. Nanochips were placed on 35-mm glass-bottom dishes (Cellvis D35-20-1-N) with the nanopillar side facing down for 100x (1.40 NA) magnification single image capturing. For 60x magnification time lapse imaging, as shown in Supplementary Video 3, nanopillars were fabricated on thin nanochips of 0.15 mm in thickness and these chips were directly imaged from their bottom. Confocal microscopy images and expansion microscope images were taken under a Nikon A1R confocal microscope equipped with 405, 488, 561, and 633 nm lasers for excitation under a 40x water objective (WD=0.61 mm, 1.15 NA). Z-stack images were taken every 0.3 µm with 1.2 Airy Unit. All other fluorescence images were taken under a Leica Thunder DMi8 System equipped with an AFC incl. closed loop focus, Leica LED8 light source, Leica filter cube set LED8, Leica external filter wheel for DMi8 and Leica K8 Camera.

### Pearson’s Correlation Coefficient Analysis

Cells were co-transfected with the indicated protein pairs, with each labeled with GFP or mCherry. Pearson’s correlation coefficient analysis was performed using ImageJ software. A 200-pixel rolling ball background subtraction was applied to both channels to subtract the cell shape-caused intensity difference. Then the cell was cropped out along the cell boundary with the nucleus region removed manually. The Pearson’s correlation coefficient (PCC) between the intensities from two channels of the cropped cell was calculated. The mean and standard deviation (STD) of the PCC was calculated and plotted.

### ER Calcium Depletion Assay

For the STIM1 and ORAI1 translocation experiments shown in Fig.4 and Supplementary Video 3, cells were transfected with the indicated probes and seeded on thin nanochips 1 day before imaging. During imaging, the cells were first imaged in Ringer’s solution, and a final concentration of 2 µM thapsigargin was administered to the imaging buffer, and the images were taken at 1 frame/second for 10 min. We selected the ORAI1 dots that were newly assembled after thapsigargin treatment and were not laterally moving, indicative of a contact localization, instead of intracellular vesicles for the kinetic analysis in Fig. 4c,d.

### Nanopillar/Nanobar Averaging and End-to-Side Ratio Quantification

Nanopillars/Nanobars were automatically identified and averaged within each cell using an in-house custom MATLAB code, as previously described^86^. Multiple cell average images (Fig 1e, 1i, 3g etc.) were generated by normalizing averaged nanopillar/nanobar images from different cells into the same dynamic scale and subsequently averaging these normalized images using ImageJ. The nanobar end-to-side ratio was determined from the averaged images of nanobars in each cell by dividing the end intensity with the side intensity, both of which were background-subtracted intensities. For ER-PM tethering protein quantification, the cells with too low of an expression level (intensity with less than 2x of the background intensity) or cells with too high of an expression level (patch size with diameter larger than 1 micron) were eliminated, and only those with intermediate intensity expression level were quantified. For all constructs in the right graph of Fig. 5l, namely all JPH3 constructs without TM domain and CAAX in this graph, a 50-pixel rolling-ball was applied to the images before quantification to subtract the cytosolic background.

### Protein Interactome Data Analysis

The protein interactome of JPH2 in living cardiomyocytes, as identified by the proximity labeling method in a prior study by Feng et al. (2020)^47^, was subjected to analysis using Metascape.org. The Biological Process (GO), Protein Function (Protein Atlas), and Subcellular Location (Protein Atlas) of these proteins were annotated. Proteins identified as associating with the plasma membrane (PM) or endosomes were grouped as PM-associated proteins. The analysis revealed that 38 proteins exhibited a prominent localization to the PM, 30 proteins displayed an additional PM localization including 2 with endosome associations, and another 3 were shown to associate with endosomes although not with the PM. Subsequently, we conducted manual assessments and identified 7 of these proteins with established roles in sensing or generating PM curvatures (Supplementary **Table1**). The expression levels of these genes in U2OS cells were assessed by the transcripts per kilobase million (TPM) value from RNA sequencing data in prior research^32^. Genes with expression levels higher but not lower than 15 TPM were knocked down. Although EHD1 was not identified in the list, we knocked down EHD1 together with EHD2 and EHD4 to avoid its complementary effects to EHD2 and EHD4.

### Statistics and Reproducibility

Statistical analysis was conducted using Prism 9 software (GraphPad). Normality in each sample group was assessed with the D’Agostino-Pearson normality test (α = 0.05). If any group within a graph did not pass the normality test, we considered this dataset to be nonparametric. For datasets that passed the normality test (i.e., parametric datasets), we employed an unpaired two-tailed Welch’s t test to evaluate the significance of the difference between two groups, and Brown-Forsythe and Welch ANOVA tests corrected with Dunnett’s T3 multiple comparisons test to compare more than two groups. For nonparametric datasets, we used the two-tailed Mann Whitney test for comparing two groups, and the Kruskal Wallis test corrected with Dunn’s multiple-comparison test for comparing more than two groups. Exact p-values are provided in the figure captions, with the significance threshold set at P < 0.05. All experiments were independently replicated at least two times, with a minimum of 3 cells per replication. Specific sample sizes (N numbers) are indicated in the figure captions.

## Data availability

All data supporting the findings of this study are available from the corresponding author upon reasonable request.

## Code availability

Customized Matlab code used in this study for nanopillar/nanobar imaging analysis was published in previous research by our lab^24^.

## Acknowledgments

We thank Dr. Richard Lewis and Dr. Ruoyi Qiu from Stanford University for insightful discussions and for sharing STIM1 and ORAI1 plasmids; Dr. Carolyn Bertozzi from Stanford University for sharing the confocal microscope; and Dr. Gea-Ny Tseng from Virginia Commonwealth University for sharing JPH2 plasmids. This study was supported by National Institutes of Health grants R35GM141598, R01HL165491, 1R01NS121934, R01 HL150693, R01 HL145676, R01 HL113006, and R01 GM144479. This work was also supported by the Stanford Sarafan ChEM-H Chemistry/Biology Interface Program 1T32GM139791-01A1 (to L.A.V.), Stanford University Center for Molecular Analysis and Design fellowship (to C-H.L.), National Institutes of Health grant Biotechnology Training Grant fellowship (to M.L.N.), American Heart Association’s Second Century Early Faculty Independence Award (to C.L.). J.L. is a Sowell Family Scholar in Medical Research. The nanofabrication in this work was performed at the Stanford Nanofabrication Facility under the NSF National Nanotechnology Coordinated Infrastructure program and Stanford Nano Shared Facilities supported by the National Science Foundation under award ECCS-2026822.

## Author Contributions

Y.Y. and B.C. conceived the study and designed the experiments. Y.Y. and L.A.V. performed the cell experiments and imaging experiments. Y.Y., L.A.V, W.Z., and W.-R.L constructed the plasmids. C.-H.L. performed the immunoprecipitation experiment. M.L.N. and L.A.V. performed the expansion microscopy experiment. C.-T.T. and Z.J. fabricated the nanostructures and performed the SEM characterization. C.L.and H.Y. performed the iPSC-CM differentiation experiments. F.S. conducted the FIB-SEM experiments. Y.Y., B.C., J.L., and J.C.W. discussed the results and wrote the manuscript with feedback from all authors.

---

**Extended Data Fig. 1:**
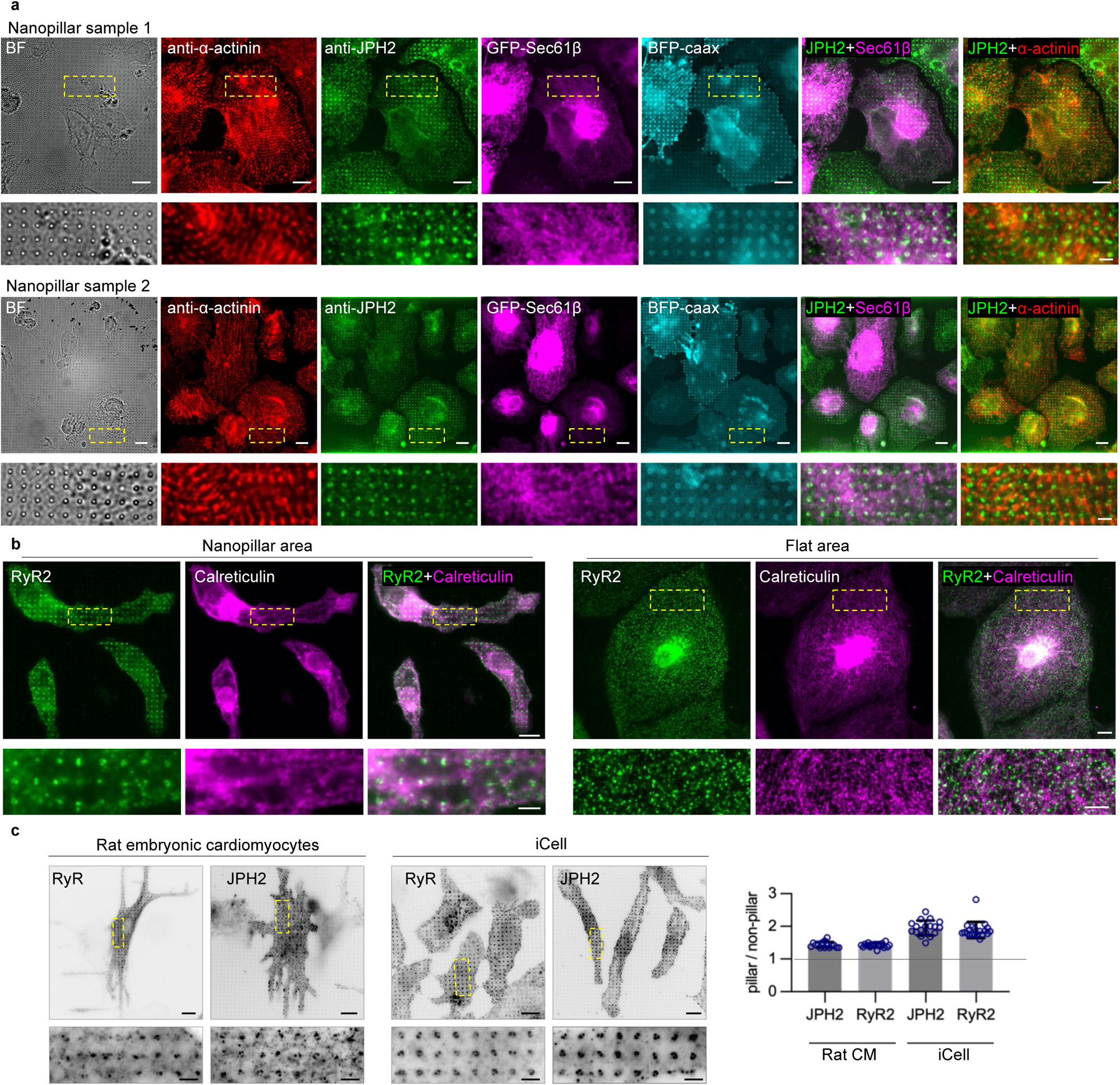
Nanopillar-induced membrane invaginations recruit dyad components in cardiomyocytes. **a**, Two additional sets of representative images of co-Immunostained JPH2 (green) and α-actinin (red) in iPSC-CMs expressing GFP-Sec61β (magenta) and BFP-CAAX (cyan) on nanopillars. The experimental condition is the same as in Fig 1d. **b**, Co-Immunostained RyR2 (green) and Calreticulin (magenta) in iPSC-CMs on nanopillar area (left) and on flat area (right). Magnified images of the yellow boxes are shown at the bottom. **c**, Immunostaining of RyR2 or JPH2 in rat embryonic CMs (left) and iCells (right). Rat embryonic CMs were dissected from E18 rat embryos, plated on nanopillars right after dissection, and cultured for 7 days before fixation and staining. iCells are human iPSC derived CMs from a commercial source (Fujifilm Cellular Dynamics). iCells were seeded on nanopillars and cultured for 51 days before staining. Inverted lookup table is used for clarity. Quantification on the right: the ratio of the fluorescent intensity at the nanopillars over the intensity in between the nanopillars quantified the same way as in Fig. 1g. n = 16, 17, 18, 20 cells for each conditions from left to right. Scale bar 10 µm in top whole cell images, 2.5 µm in bottom zoom-in images for (**a**,**b**,**c**). All experiments were independently replicated at least two times. All error bars represent STD.

**Extended Data Fig. 2:**
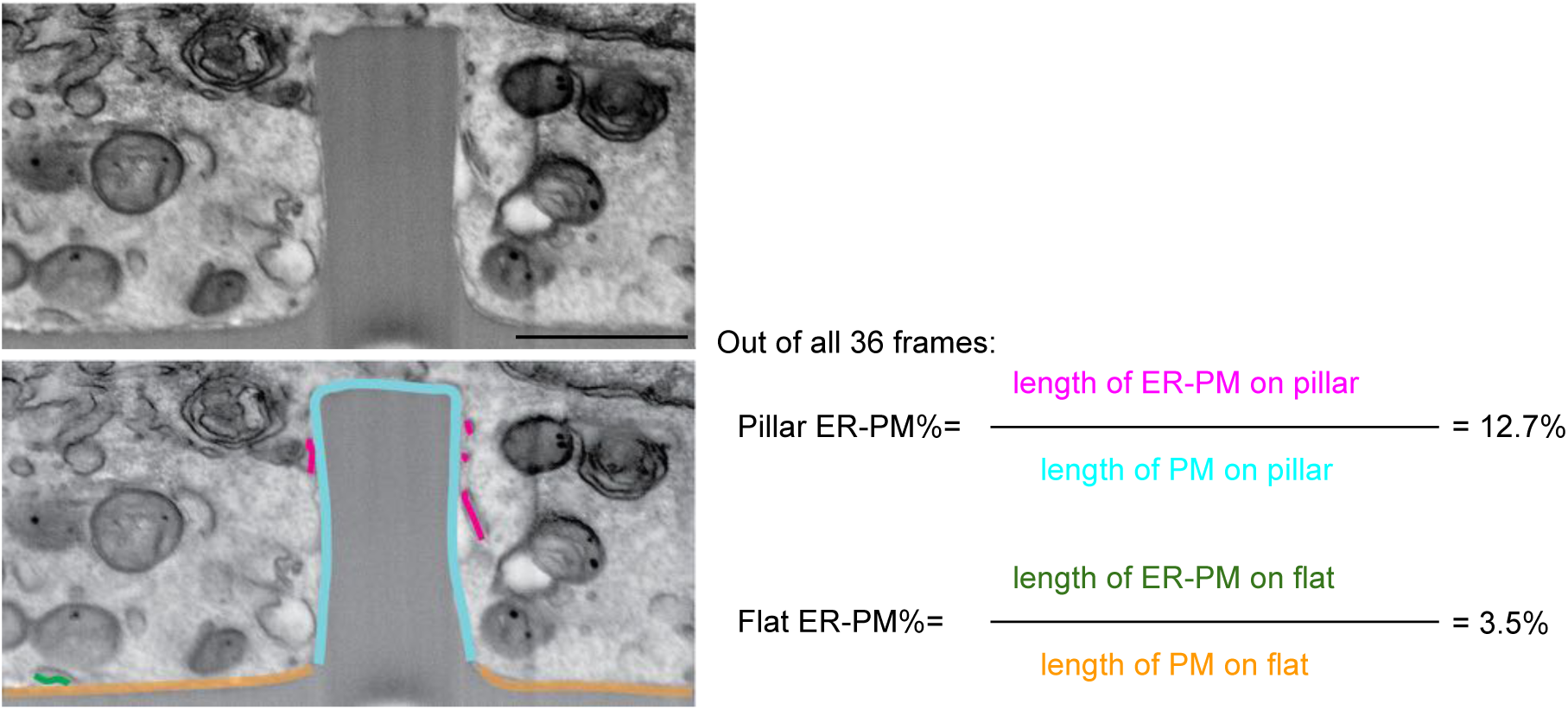
Illustration of the method to quantify the ER-PM contact overage in FIB-SEM images. One section of the FIB-SEM images shown in Fig. 2c is shown as an example. Upper image is the original image, and the lower image is the same image with the Length of the ER-PM contact on pillar (magenta), the ER-PM contact on the flat (green), the total length of membrane on the pillar (cyan), and the total length of membrane on the flat (orange) color-highlighted. Data from 36 distinct frames were manually collected and the ER-PM contact coverage on nanopillar area and on flat area were then calculated respectively as indicated on the right. Scale bar 100 nm. The experiment was independently replicated two times.

**Extended Data Fig. 3:**
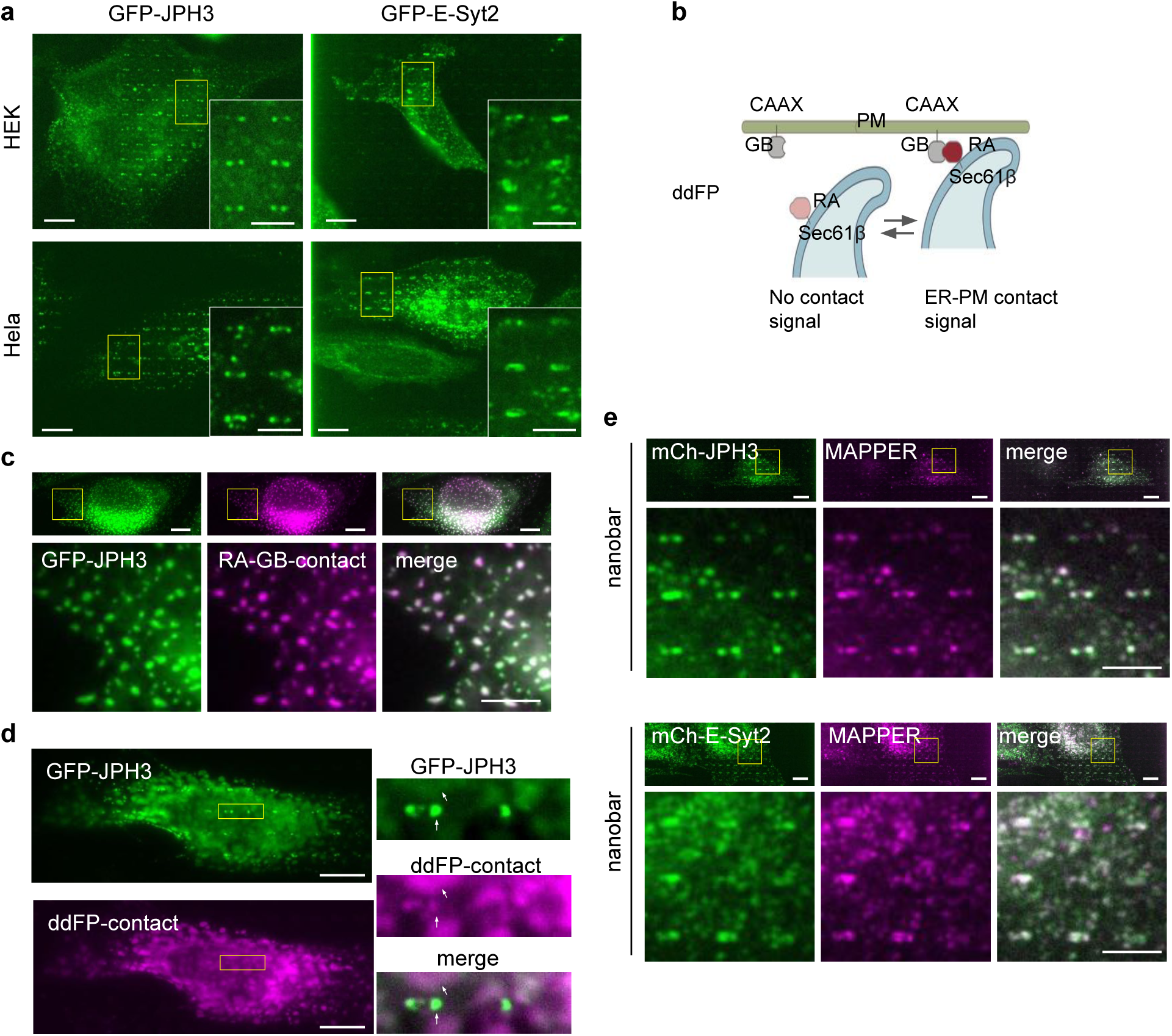
Distinct spatial distribution of JPH3 and E-Syt2. **a**, Representative images of GFP-JPH3 and GFP-E-Syt2 expressed HEK-297T cells and Hela cells. The quantifications are included in Fig. 3h. Insets show the enlarged 6-nanbar regions in yellow boxes. **b**, A schematic illustration of the ddFP-based ER-PM contact sensor. **c**, ddFP-based ER-PM contact sensor (magenta) and co-expressed GFP-JPH3 (left, green) show extensive colocalization in U2OS cells cultured on flat surfaces. Regions in the yellow boxes are enlarged in the bottom row. **d**, On nanobars, GFP-JPH3 shows stronger preference toward nanobar ends than ddFP-based ER-PM contact sensors. Two-bar region in yellow boxes were enlarged on the right. White arrows point to two distinct contact sites at the nanobar end and the flat surface. **e**, Representative images of GFP-MAPPER (magenta) co-expressed with mCherry-JPH3 (top, green) or mCherry-E-Syt2 (bottom, green) on nanobars. Region in the yellow boxes are enlarged in the bottom row. Scale bar 10 µm in whole cell images, 5 µm in zoom-in images for (**a**,**c**,**d**,**e**). All experiments were independently replicated at least three times.

**Extended Data Fig. 4:**
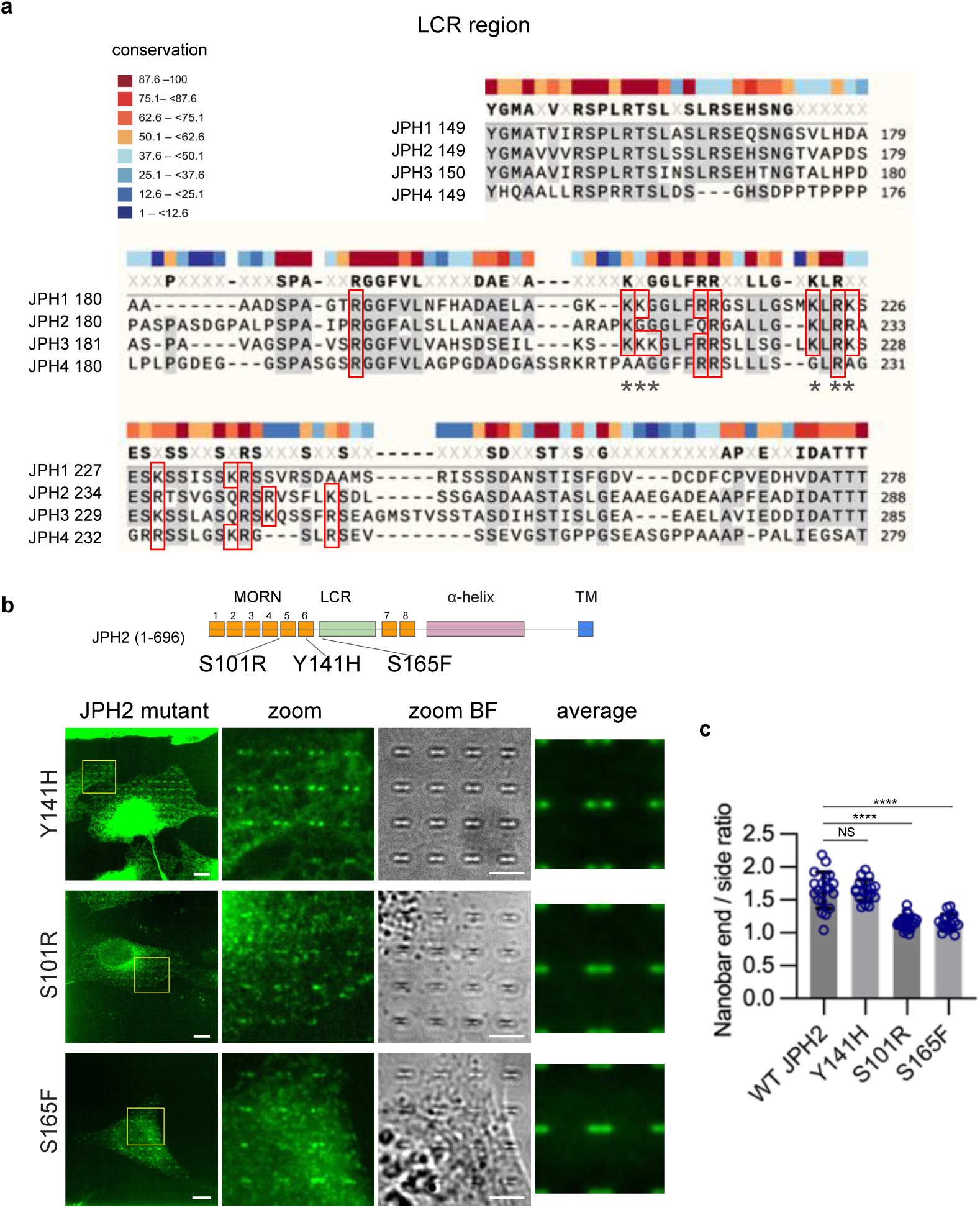
LCRs for the four human JPH genes show conserved polybasic residues, and hypertrophic cardiomyopathy associated point mutations impair membrane curvature targeting of JPH2. **a**, Alignments are generated from Snapgene. The positively charged residues are contained in the red boxes. Conserved residues and substitutions are colored as indicated. The numbers on the left refer to the position of the first residue shown in each row. The * marked amino acids were mutated into alanine (K210A, K211A, K212A, K224A, R226A, K227A) in the LCR_KRtoA mutant in Fig. 5e. **b**, Representative images of U2OS cells expressing GFP-tagged JPH2 mutants associated with hypertrophic cardiomyopathy. Left: whole cell images; Middle: enlarged fluorescent view and bright field view of the regions in yellow boxes; Right: averaged nanobar images from multiple cells. Cell number: n = 19 (Y141H), 25 (S101R), 18 (S165F). Diagram of the position of each point mutation shown on the top. Scale bar 10 µm in whole cell images, 5 µm in zoom-in images. **c**, Quantifications of the nanobar end-to-side ratios for WT-JPH2, JPH2-Y141H, JPH2-S101R, and JPH2-S165F. Cell number is the same as in (**b**). ****P < 0.0001, NS P>0.9999. All experiments were independently replicated at least three times. All error bars represent STD. Brown-Forsythe and Welch ANOVA tests was used to assess significance for (**c**).

**Extended Data Fig. 5:**
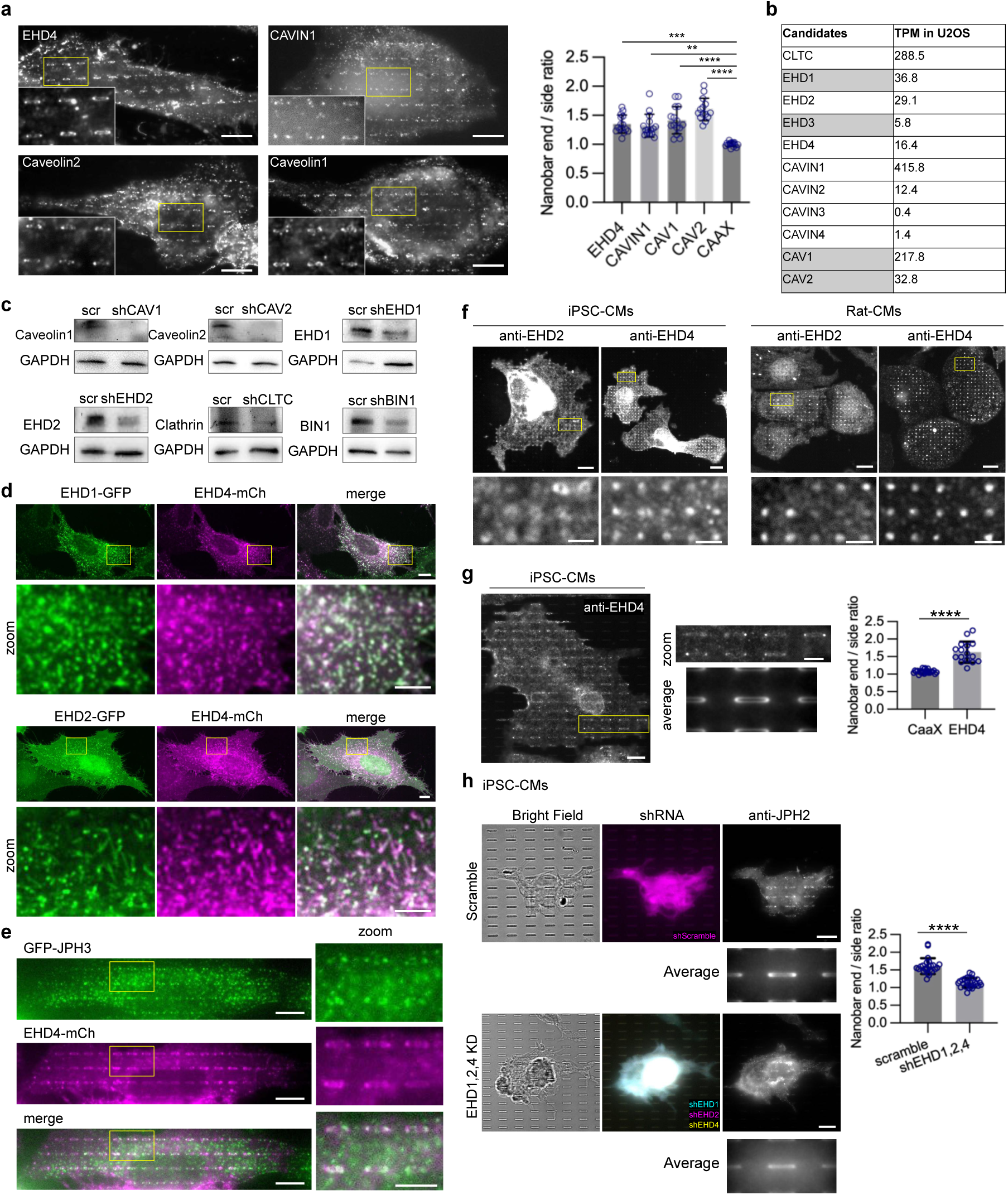
EHDs sense PM curvatures and play important roles in JPH’s membrane curvature targeting in both U2OS cells and iPSC-CMs. **a**, EHD4, CAVIN1, Caveolin1, and Caveolin2 all show preferential accumulations at the nanobar ends. Quantifications of the end-to-side ratios are shown on the right. Cell number: n =14 cells for Caveolin2, n = 15 cells for all other groups. ***P=0.0001, **P=0.0015; ****P<0.0001. **b**, mRNA transcript levels in U2OS cells (shown as normalized transcripts per million, nTPM), obtained from protein atlas database. Genes highlighted in gray: not identified in the JPH2 interactome but are closely related either genetically or functionally. Genes including EHD3, CAVIN2, CAVIN3, and CAVIN4 were not selected as knockdown targets in KD experiments because of their low TPM level in U2OS cells. **c**, Western blots confirming the effective shRNA knockdown of CAV1, CAV2, EHD1, EHD2, CLTC (clathrin heavy chain), and BIN1. GAPDH was blotted as a loading control. **d**, Representative images of U2OS cells co-expressing EHD4-mCherry with EHD1-GFP (Top) or with EHD2-GFP (Bottom). Enlarged views are zoomed from the yellow boxes in the whole cell images. **e**, Representative images of U2OS cells expressing GFP-JPH3 (green) and EHD4-mCh (magenta) seeding on nanobars. Enlarged views are zoomed from the yellow boxes in whole cell images. **f**, Representative images of immunostaining of EHD2 or EHD4 in iPSC-CMs or rat embryonic CMs on nanopillars. Magnified images of the region in yellow boxes are shown in the bottom row. Scale bar 10 µm in the top row, 2.5 µm in the bottom row. **g**, Representative images of immunostaining of EHD4 in iPSC-CMs on nanobars. Magnified images of the region in yellow boxes are shown in the middle row. Averaged fluorescent signals of all nanobars from multiple cells were averaged and displayed in the bottom row. n = 16 cells for the average image. Bottom graph: quantification of the end to side ratios of EHD4 compared to that of BFP-CAAX expressed in iPSC-CM on nanobars. Cell number: CAAX = 23 (same data as in Fig.1k), EHD4 = 16. ****P <0.0001. **h**, Representative images of immunostaining of JPH2 in iPSC cells transfected with shRNA of scramble (top) or EHD1/2/4 (bottom). From left to right: Bright field image, fluorescent protein expression indicating successful transfection of shRNA, and JPH2 staining. Averaged nanobar images of JPH2 staining from multiple cells are shown at the bottom of JPH staining panels. Quantifications of the end-to-side ratios are shown on the right. Cell number for both average and quantification: Scramble = 24, shEHD1/2/4 = 26. ****P <0.0001. Scale bars 10 µm in all whole cell images, 5 µm in all zoomed images unless otherwise mentioned. All experiments were independently replicated at least two times. All error bars represent STD. Kruskal–Wallis test corrected with Dunn’s multiple-comparison test was used to assess significance in (**a**). Unpaired two-tailed Welch’s t test was used to assess significance in (**g**). Two-tailed Mann-Whitney test was used to assess significance in (**h**).

**Extended Data Fig. 6:**
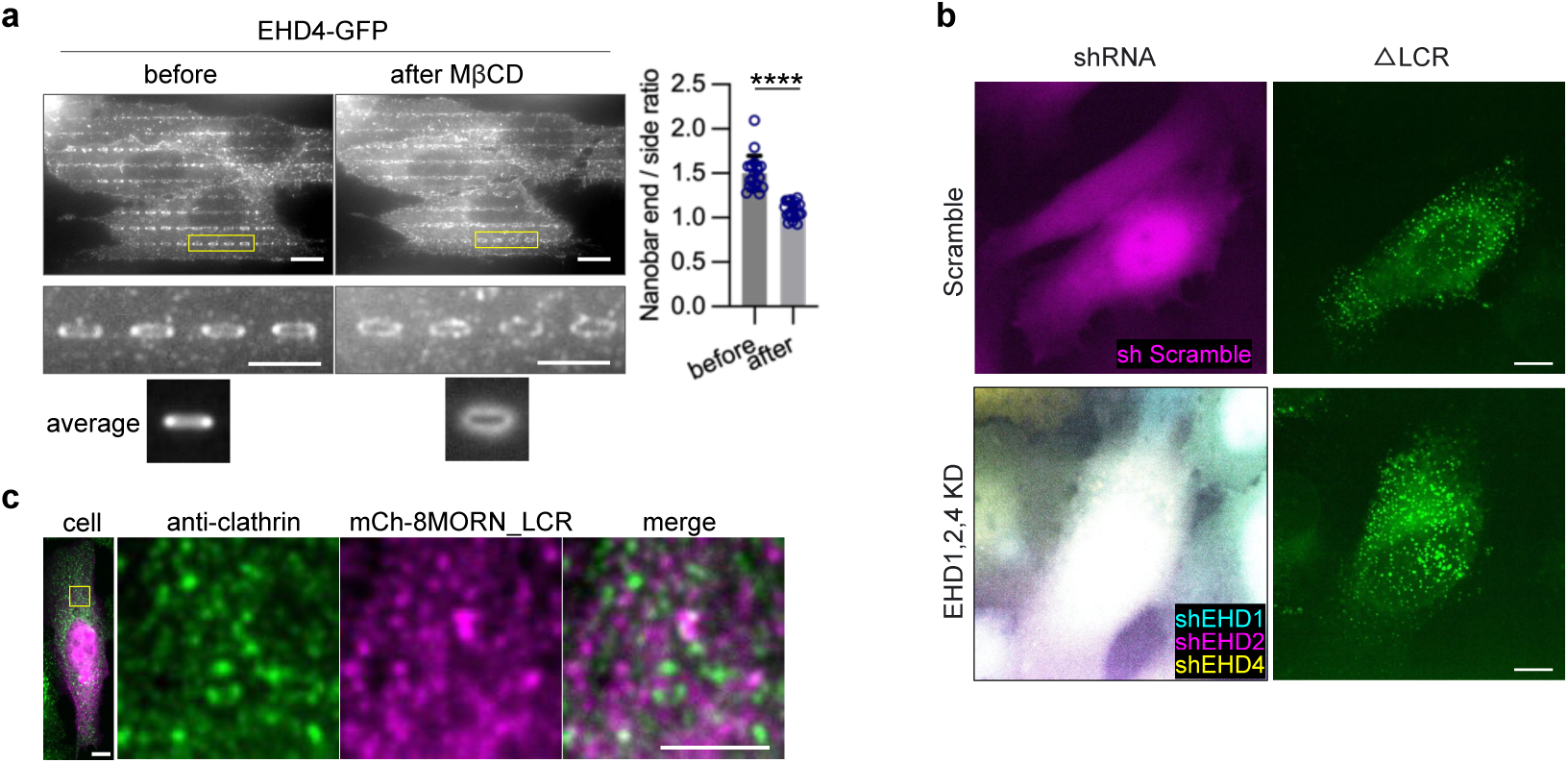
EHD4’s PM curvature sensitivity is cholesterol dependent, MORN motifs possess PM affinity and MORN_LCR interacts with EHD4. **a**, The distribution of EHD4-GFP on nanobars before and after 10 mM MβCD treatment at 37°C for 30 min. Magnified images of the yellow boxes are shown in the middle row. Averaged fluorescent signals of all nanobars from multiple cells were averaged and displayed in the bottom row. Quantifications of the end-to-side ratios are shown on the right. n = 15 cells for both conditions. ****P <0.0001. **b**, Representative images of GFP-ΔLCR expressed in EHD-1,-2,-4 triple knockdown U2OS cells (bottom) or scramble knockdown control U2OS cells (top). **c**, Representative images of U2OS cells expressing mCherry-8MORN_LCR (magenta) and immunostained with clathrin-heavy-chain antibody (green). Enlarged views are zoomed from the yellow boxes in whole cell images. Quantification of this data was shown in Fig.6h. Scale bar 10 µm in all whole cell images, 5 µm in all zoomed images. All experiments were independently replicated at least two times. All error bars represent STD. Unpaired two-tailed Welch’s t test was used to assess the significance in (**a**).

**Extended Data Fig. 7:**
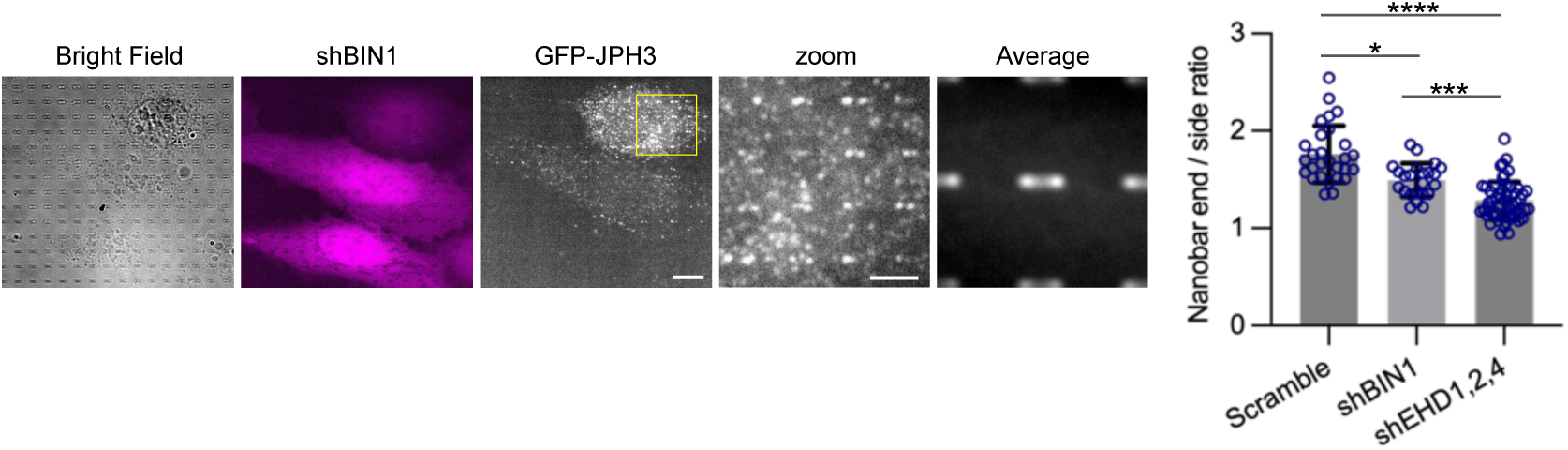
BIN1 knockdown mildly reduces the curvature preference of JPH3. Representative images of GFP-JPH3 in U2OS expressing a BIN1 shRNA. Bright field and the fluorescent protein co-expressed with shRNA are shown on the left. Zoom is the enlarged view of the region in yellow box. Averaged image of all nanobars from multiple cells is shown on the right. Cell number for average: Scramble = 28; shBIN1 = 23; shEHD1,2,4 = 55. Scale bars all 10 µm in whole cell, 5 µm in zoomed image. The experiment was independently replicated for three times. All error bars represent STD. Kruskal–Wallis test corrected with Dunn’s multiple-comparison test was used to assess significance. *P=0.0382, ***P=0.0008, ****P<0.0001.

## Supplementary Video Cations

Supplementary Video1:

**FIB-SEM image stack of ER-PM contact from HL-1 cells growing on nanopillars.** FIB-SEM image stacks of the structure shown in Fig. 2c. ER forms multiple contact sites with the curved PM at the nanopillar. Scale bar 1 µm.

Supplementary Video2:

**Three-dimensional animation of ExM imaging of the cardiomyocyte seeding on nanopillars from** Fig. 2d. GFP-CAAX in green and anti-JPH2 in magenta. Scale after expansion is indicated on the grids.

Supplementary Video3:

**Timelapse of ORAI1 and STIM1 in response to calcium store depletion at membrane curvatures.** U2OS cells were co-transfected with ORAI1-GFP (left) and mCherry-STIM1(right) and plated on nanobar chips 24 hours before imaging. During recording, 2 µM Thapsigargin (Tg) was added to induce calcium store depletion at the 4^th^ frame. Frame interval was 3s. The video is played at a speed 75x faster than real time. Scale bar 10 µm.

## Notes

### Competing Interest Statement

The authors have declared no competing interest.

