## Supplementary Table S1 for "Membrane Curvature Promotes ER-PM Contact Formation via Junctophilin-EHD Interactions"

Plasma membrane (Approved, Enhanced, Supported)

| Majority_prot | Peptide_count | Peptide_count | Peptide_count | Protein_name | Gene_names |
|---|---|---|---|---|---|
| Q9Z1W8 | 4 | 1 | 1 | Potassium-tra | Atp12a |
| P53986 | 3 | 3 | 3 | Monocarboxy | Slc16a1 |
| P26231 | 4 | 4 | 3 | Catenin alpha | Ctnna1 |
| Q9ES83 | 18 | 18 | 18 | Blood vessel | Bves |
| P48193 | 2 | 2 | 2 | Protein 4.1 | Epb41 |
| Q02257 | 9 | 9 | 9 | Junction plak | Jup |
| E9Q9K5 | 1 | 1 | 1 | Triadin | Trdn |
| P14069 | 1 | 1 | 1 | Protein S100-A | S100a6 |
| P14602 | 11 | 11 | 11 | Heat shock pro | Hspb1 |
| P17182 | 2 | 2 | 2 | Alpha-enolase | Eno1 |
| P21550 | 2 | 2 | 2 | Beta-enolase | Eno3 |
| P23927 | 19 | 19 | 19 | Alpha-crystal | Cryab |
| O88492 | 35 | 35 | 35 | Perilipin-4 | Plin4 |
| P14206 | 3 | 3 | 3 | 40S ribosomal | Rpsa |
| Q99JB2 | 1 | 1 | 1 | Stomatin-like | Stoml2 |
| Q9QZF2 | 1 | 1 | 1 | Glypican-1;S | Gpc1 |
| Q3UHA3 | 1 | 1 | 1 | Spatacsin | Spg11 |
| P09055 | 2 | 2 | 2 | Integrin beta- | Itgb1 |
| **A2AMM0** | **9** | **9** | **9** | **Muscle-relat** | **Murc** |
| **Q91VJ2** | **2** | **2** | **2** | **Protein kina** | **Prkcdbp** |
| **Q9EQP2** | **3** | **3** | **3** | **EH domain-c** | **Ehd4** |
| Q9QUM0 | 1 | 1 | 1 | Integrin alpha | Itga2b |
| Q9R0P5 | 1 | 1 | 1 | Destrin | Dstn |
| P07356 | 5 | 5 | 5 | Annexin A2 | Anxa2 |
| Q60714 | 10 | 10 | 10 | Long-chain fa | Slc27a1 |
| Q8CI51 | 7 | 7 | 7 | PDZ and LIM | Pdlim5 |
| Q8BGK2 | 4 | 4 | 4 | [Protein ADP | Adprhl1 |
| P59913 | 7 | 7 | 7 | Protein-L-iso | Pcmtd1 |
| **Q63918** | **11** | **11** | **11** | **Serum depri** | **Sdpr** |
| E9Q6P5 | 2 | 2 | 2 |  | Ttc7b |
| G5E8K5 | 9 | 9 | 6 | Ankyrin-3 | Ank3 |
| **Q8BH64** | **2** | **2** | **2** | **EH domain-c** | **Ehd2** |
| Q9WTR5 | 2 | 2 | 2 | Cadherin-13 | Cdh13 |
| Q9Z2C5 | 26 | 26 | 26 | Myotubularin | Mtm1 |
| P15116 | 1 | 1 | 1 | Cadherin-2 | Cdh2 |
| Q80UL9 | 1 | 1 | 1 | Junctional ad | Amica1 |
| **O54724** | **17** | **17** | **17** | **Polymerase** | **Ptrf** |
| Q62165 | 4 | 4 | 4 | Dystroglycan | Dag1 |

Plasma membrane (Additional)

| Majority_prot | Peptide_count | Peptide_count | Peptide_count | Protein_name | Gene_names |
|---|---|---|---|---|---|
| A2AUC9 | 1 | 1 | 1 | Kelch-like pro | Klhl41 |

| | | | | | |
|---|---|---|---|---|---|
| P97447 | 2 | 2 | 2 | Four and a ha | Fhl1 |
| P83882 | 3 | 3 | 3 | 60S ribosoma | Rpl36a |
| D3YXG0 | 2 | 2 | 2 | | Hmcn1 |
| P27546 | 2 | 2 | 2 | Microtubule-a | Map4 |
| P63254 | 3 | 3 | 3 | Cysteine-rich | Crip1 |
| Q9Z239 | 1 | 1 | 1 | Phospholemr | Fxyd1 |
| Q9JKV1 | 1 | 1 | 1 | Proteasomal | Adrm1 |
| E9Q401 | 392 | 392 | 364 | Ryanodine re | Ryr2 |
| Q3U7R1 | 2 | 2 | 2 | Extended syr | Esyt1 |
| P97414 | 7 | 7 | 7 | Potassium vo | Kcnq1 |
| P35278 | 1 | 1 | 1 | Ras-related p | Rab5c |
| Q9WUM5 | 4 | 4 | 4 | Succinyl-CoA | Suclg1 |
| P63250 | 7 | 7 | 7 | G protein-act | Kcnj3 |
| Q8R035 | 1 | 1 | 1 | Peptidyl-tRNA | Ict1 |
| Q9CQA3 | 16 | 16 | 16 | Succinate de | Sdhb |
| Q3TL44 | 7 | 7 | 7 | NLR family m | Nlrx1 |
| A2ARV4 | 1 | 1 | 1 | Low-density l | Lrp2 |
| O35682 | 1 | 1 | 1 | Myeloid-asso | Myadm |
| Q9DCT8 | 10 | 10 | 10 | Cysteine-rich | Crip2 |
| Q8BKZ9 | 5 | 5 | 5 | Pyruvate deh | Pdhx |
| Q5FW52 | 2 | 2 | 2 | Muscular LMI | Mlip |
| P70414 | 7 | 7 | 7 | Sodium/calciu | Slc8a1 |
| Q1XH17 | 6 | 6 | 6 | Tripartite mot | Trim72 |
| P61021 | 1 | 1 | 1 | Ras-related p | Rab5b |
| O88952 | 2 | 2 | 2 | Protein lin-7 h | Lin7c |
| P05125 | 3 | 3 | 3 | Natriuretic pe | Nppa |
| P14094 | 10 | 10 | 10 | Sodium/potas | Atp1b1 |
| Q924D0 | 2 | 2 | 2 | Reticulon-4-ir | Rtn4ip1 |
| Q9WV27 | 8 | 1 | 1 | Sodium/potas | Atp1a4 |

Endosome

| Majority_prot | Peptide_count | Peptide_count | Peptide_count | Protein_name | Gene_names |
|---|---|---|---|---|---|
| **Q68FD5** | **2** | **2** | **2** | **Clathrin hea** | **Cltc** |
| Q91V41 | 1 | 1 | 1 | Ras-related p | Rab14 |
| P35282 | 1 | 1 | 1 | Ras-related p | Rab21 |

Plasma membrane and curvature

| Majority_prot | Peptide_count | Peptide_count | Peptide_count | Protein_name | Gene_names |
|---|---|---|---|---|---|
| **A2AMM0** | **9** | **9** | **9** | **Muscle-relat** | **Murc** |
| **Q91VJ2** | **2** | **2** | **2** | **Protein kina** | **Prkcdbp** |
| **Q9EQP2** | **3** | **3** | **3** | **EH domain-c** | **Ehd4** |
| **Q63918** | **11** | **11** | **11** | **Serum depri** | **Sdpr** |

| | | | | | |
|---|---|---|---|---|---|
| Q8BH64 | 2 | 2 | 2 | EH domain-c | Ehd2 |
| O54724 | 17 | 17 | 17 | Polymerase | Ptrf |
| Q68FD5 | 2 | 2 | 2 | Clathrin hea | Cltc |
